# Petagraph: A large-scale unifying knowledge graph framework for integrating biomolecular and biomedical data

**DOI:** 10.1101/2023.02.11.528088

**Authors:** Benjamin J. Stear, Taha Mohseni Ahooyi, Shubha Vasisht, Alan Simmons, Katherine Beigel, Tiffany J. Callahan, Jonathan C. Silverstein, Deanne M. Taylor

## Abstract

The use of biomedical knowledge graphs (BMKG) for knowledge representation and data integration has increased drastically in the past several years due to the size, diversity, and complexity of biomedical datasets and databases. Data extraction from a single dataset or database is usually not particularly challenging. However, if a scientific question must rely on integrative analysis across multiple databases or datasets, it can often take many hours to correctly and reproducibly extract and integrate data towards effective analysis. To overcome this issue, we created Petagraph, a large-scale BMKG that integrates biomolecular data into a schema incorporating the Unified Medical Language System (UMLS). Petagraph is instantiated on the Neo4j graph platform, and to date, has fifteen integrated biomolecular datasets. The majority of the data consists of entities or relationships related to genes, animal models, human phenotypes, drugs, and chemicals. Quantitative data sets containing values from gene expression analyses, chromatin organization, and genetic analyses have also been included. By incorporating models of biomolecular data types, the datasets can be traversed with hundreds of ontologies and controlled vocabularies native to the UMLS, effectively bringing the data to the ontologies. Petagraph allows users to analyze relationships between complex multi-omics data quickly and efficiently.

## Introduction

As biological datasets continue to grow in size and complexity, knowledge graphs and machine learning approaches applied to knowledge graphs are likely the best near-term solutions for integrating across large and disparate biomedical datasets (Nicholson and Greene 2020). Knowledge graphs are graph-structured data collections that integrate data points and their relationships into a connected graph entity. Knowledge graphs have been used for many applications, including drug discovery and repurposing, target detection, and prediction (Alshahrani and Hoehndorf 2018; Moon et al. 2021; Zheng et al. 2021; Alves et al. 2021). Other applications include integration and analysis of heterogeneous COVID-19 data (Steenwinckel et al. 2020; Cernile et al. 2021; Reese et al. 2021; Domingo-Fernández et al. 2021; Zhang et al. 2021; Ostaszewski et al. 2021; Chen et al. 2021), oncology research (Zhu et al. 2023; Zhao et al. 2023; Jha et al.), and gene-disease associations (Choi and Lee 2021; Alves et al. 2021) among many others. The unifying intent in these projects is the meaningful integration of heterogeneous biological data for exploration, analysis, and discovery. Machine-learning approaches can also work with knowledge graphs, including graph-specific methods such as GNNs (Wu et al. 2021); (Kipf and Welling 2016).

Creating biomedical knowledge graphs from multi-omic biomolecular data can involve significant expenditures in human, computational, and storage resources. Still, a knowledge graph can offer significant advantages to research efforts once the graph is constructed, including providing a rich, connected environment for exploration and modeling. We thus constructed Petagraph, a knowledge graph framework based on the Unified Biomedical Knowledge Graph (UBKG) (Silverstein et al. 2023). The UBKG is a property graph representation of the NIH Unified Medical Language Service (UMLS) (Bodenreider 2004), enabling Petagraph to extend the UMLS data model. Petagraph essentially supports integrating large-scale biomolecular and biomedical data types into a UBKG environment of about 200 cross-referenced ontologies. Thus, Petagraph’s model provides a rich environment where relationships can be easily established between heterogeneous data types, more richly connecting the graph.

Petagraph was conceived as a knowledge graph for rapid feature selection to explore candidates for gene variant epistasis. The graph was assembled on the UBKG scaffold with new models for quantitative data from RNA-seq, variant, and model organism datasets from projects such as GTEx (Lonsdale et al. 2013), HuBMAP (HuBMAP Consortium 2019), Kids First (Gabriella Miller Kids First Pediatric Research Program (Kids First)-The Office of Strategic Coordination-The Common Fund – National Institutes of Health), and IMPC (Groza et al. 2023). We summarized these and other datasets to generate valuable biological assertions with reduced dimensions (compared to the original data), which still retain information about the biological processes they represent. For example, we calculated the median TPM value for each gene for each GTEx tissue to represent a reduced-dimension representation of tissue-specific gene expression.

To facilitate the integration of these and other biomolecular datasets into the UMLS ontology schema, we also included several supporting ontologies and datasets, which will now become permanent additions to the UBKG (Table S1). The number of supporting ontologies for biomolecular data included in Petagraph also allows for customization with particular datasets for specific use cases. Petagraph’s biomolecular data model is also designed to ingest and support several different genomic data types. The latest version of Petagraph also introduces a Chromosome Region ontology to easily associate relevant features with different resolutions by chromosome position and chromosomal vicinity (**Figures S1-S2**).

New datasets are easily incorporated into the Petagraph model as outlined in the Methods section. We show that Petagraph’s integrated biomolecular data model can yield meaningful results for query-based use cases that generate rapid feature selection across disparate data types.

## METHODS

### Concepts, Codes, and Term nodes

The UBKG and thus Petagraph utilizes the UMLS data model containing Concepts, Codes, and Terms that describe specific categories of data that carry information. These categories are discussed in detail in the UMLS Metathesaurus (National Library of Medicine (US) 2009). Within Petagraph, “Concept nodes” are the central backbone of the entire graph model. The Concept node is a graph representation of a UMLS Concept and thus represents a universal hub for a particular conceptual meaning from external references. For example, a Concept node for the human gene TP53 may represent that gene’s definition or version for a chosen set of references. Every Concept node is assigned a Concept Unique Identifier (CUI) based on the UMLS model. For example, the UMLS’s CUI for the *H. sapiens* TP53 gene is ‘C0079419.’ CUIs are simply alphanumeric identifiers and are not informative outside of their use as an identifier. “Code nodes” identify the external reference IDs that connect to that Concept node. Examples of three Code nodes connecting to the Concept node for TP53 could be “HGNC:11998” (HGNC ID), “ENSG00000141510” (Ensembl), and “7157” (NCBI Gene ID). Thus a Concept can be defined by its Code nodes or Term nodes, which can act as definitions or names. A Term node can provide human-readable information about an attached Code node. In the case of TP53, the Code node “HGNC:11998” would have the Term labeled “tumor protein p53” (**Figure 1**).

**Figure 1:**
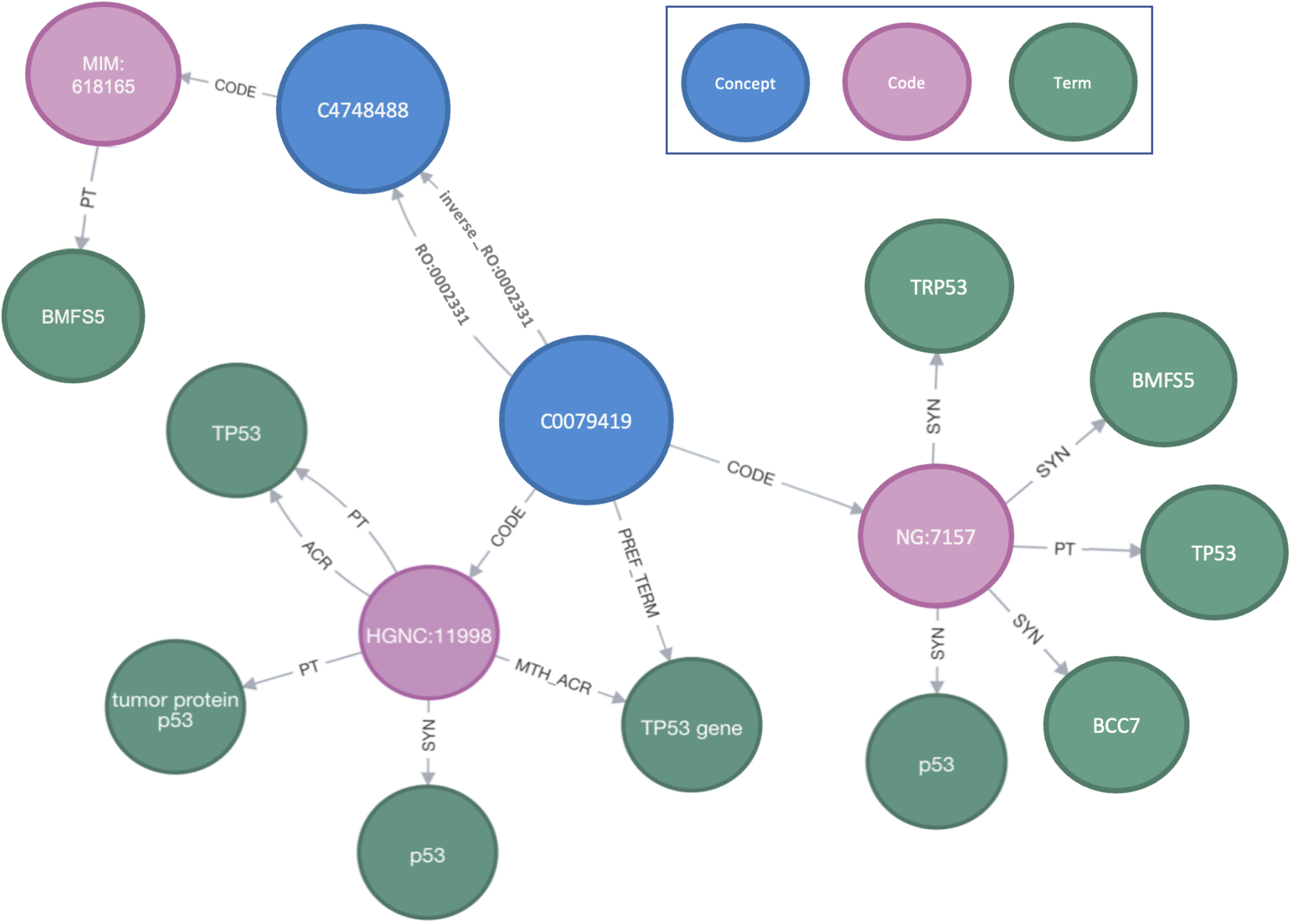
Example graph schema. Example Concept nodes (blue) connecting to associated Code nodes (pink) and Term codes (green). The Concept in blue for TP53 (C0079419) with two of its codes and associated terms are shown: HGNC (HGNC:11998) and NCBI Gene (NG:7157). Other codes also exist for C0079419 but are not shown. Note that all the relationships from the concept to codes and terms are unidirectional: directed outwards. The only bidirectional connections allowed in Petagraph are those between Concept nodes, here shown as TP53’s Concept and one of the OMIM phenotypes associated with TP53, connected by the OBO relations ontology (RO:0002331) for “involved in”. Petagraph has many millions of connected concept nodes. ACR: Abbreviation. MTH_ACR: Abbreviation. PT: Preferred. PREF_TERM: Preferred term. SYN: Synonym.

Petagraph’s data model is built around the UBKG’s property graph model of the UMLS “Concept” and “Relationship” classes with specific directionalities and is, therefore, a directed graph. A “Concept” in Petagraph’s model is a generic node class allowed to connect bidirectionally within its class and is the only node type allowed to have bidirectional edges. Concept nodes can also connect outwardly to other node classes, either “Code nodes” representing data system IDs, or “Term Nodes,” representing details such as descriptions or definitions, often connected through Code Nodes. In **Figure 1**, two Concept nodes are connected bidirectionally using specific edge types: C0079319 (TP53) and C4748488 (OMIM. Each Code term is connected to its assigned Concept node outwardly. Each Term node is connected to its Code node. Controlling the directionality of relationships (edges) in the graph supports the central role that ‘Concepts’ play in the data structure and limits the number of paths that can be traveled when integrating data types among different datasets and systems. There are hundreds of edge types in the Petagraph property graph model supporting specific relationships among Concept nodes, many of which derive from the UMLS relationship classes directly, while others are designed specifically based on the datasets ingested. A document covering aspects of the Petagraph schema can be found at: https://github.com/TaylorResearchLab/Petagraph/blob/main/schema_docs/schema_reference.md.

We will frequently elide “nodes” when referring to Concept nodes, Code nodes, and Term nodes in this manuscript for ease of reading.

### Standards, Ontologies, and supporting data

The UBKG includes, at the time of this writing, 185 ontological and classification systems, 165 of which are found in the UMLS. Petagraph additionally includes other reference and interrelated datasets to support the integration and querying of biomolecular data (**Table S1**).

### Data Ingestion Process

The Petagraph database is instantiated on a Neo4j graph database, version 4.3.

Scripts have been developed as part of the UBKG project for data ingestion and code for the UBKG data ingestion pipeline is featured in the project’s GitHub repository (Silverstein et al. 2023). The scripts support ingesting data that utilize the same identifiers (such as gene name, Human Phenotype Ontology (HP) ID, or others). In the case that a Concept is already in the UBKG, the existing CUI is utilized and the new Code and Term(s) added. In cases where we introduce ontologies or data summaries that do not have an existing CUI, the UBKG ingestion process can automatically generate a new CUI. We have made minor adaptations in the UBKG scripts for loading biomolecular data for Petagraph, and those scripts have been included in the Petagraph GitHub repository.

The data loading pipeline contains four major steps (**Figure 2**): (1) Data selection and modeling, (2) Cleaning and preparing data, (3) Executing the UBKG “OWLNETS” pythons scripts and (4) Import using the Neo4j Bulk Import tool. We discuss each of these steps in turn.

**Figure 2.**
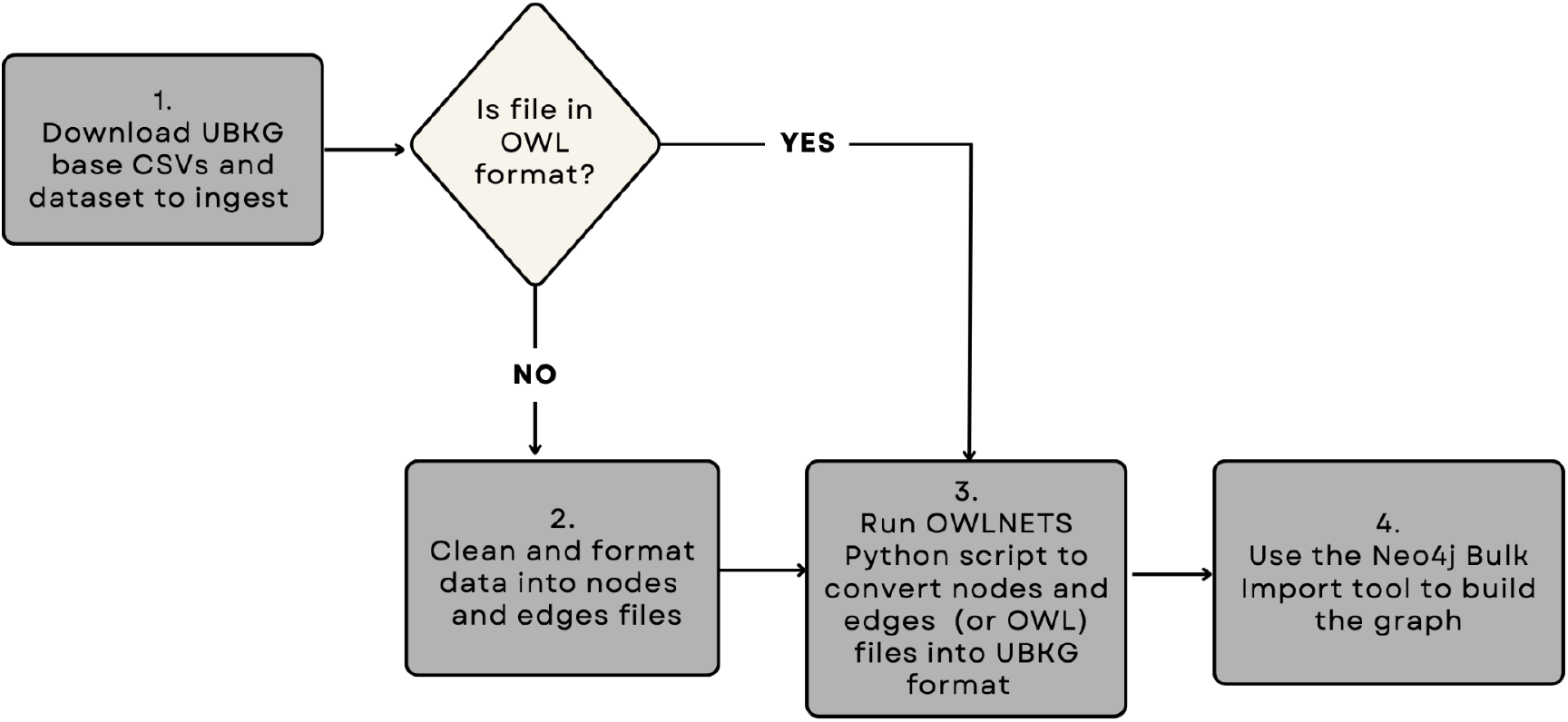
Petagraph Data Ingestion Workflow. The data ingestion workflow starts by downloading the UBKG base CSVs and the dataset(s) that will be integrated. Then the raw data (or ontology) must be formatted into nodes and edges file, the guidelines of which are in the user guide. If the file is in OWL format then no extra formatting needs to be done and this step can be skipped. The third step involves running the OWLNETS Python script which will convert the edges and nodes files into the UBKG format. Once the files are in UBKG format They are simply appended to the base UBKG files. Lastly, Neo4js command-line bulk import tool will be used to build the graph database.

#### Step 1: Data selection and modeling

The selection, sourcing, review, and modeling of data appropriate for the knowledge graph requires a comprehensive understanding of the knowledge graph’s existing or planned model and structure. When creating a model that merges disparate types of biomolecular data, there should be an emphasis on modeling the relationships that are meaningful and biologically relevant to the data. For example, when modeling relationships between Concepts for genes and Concepts for associated regulatory motifs, biological context (such as the proximity of a regulatory motif to a gene within the genome) should be considered in the model of the relationship. More complex models are used to support experimental data. Petagraph adds data models and additional data onto the UBKG by appending nodes and relationships to those in the base CSVs.

#### Step 2: Manual work in cleaning data into preliminary nodes and edges files

Clean and format the data. If the data is in OWL format then this step can be skipped and the OWL file can be run directly with the OWLNETS.py script (see https://github.com/callahantiff/PheKnowLator/wiki/OWL-NETS-2.0). However, most of the datasets we ingested are not in OWL format, nor are they proper ontologies, so a fair amount of reformatting and modeling was necessary for almost all of the datasets. After deciding exactly how a specific data set was going to be modeled, data cleaning mostly involved removing missing or irrelevant data points. Data formatting consisted of organizing the datasets into rows of ‘triples’, or node-edge-node rows. This ‘edges.tsv’ file represents all the Concept-to-Concept relationships for the given dataset. The ‘nodes.tsv’ file consists of the Code ID for the corresponding Concept nodes in the ‘edges.tsv’ file as well as various metadata fields included in the Terms for the Code nodes.

#### Step 3: Execution of the UBKG ‘OWLNETS’ python ingestion script to convert nodes and edges files into UBKG format

Download the necessary UBKG CSVs. The UBKG exists as a set of eleven ‘base’ CSV files that represent the nodes and edges of the graph. The UBKG CSVs range in size from 5KB to 1GB. Once the nodes and edges files have been created we use a python script called OWLNETS (that can be found in GitHub: (https://github.com/TaylorResearchLab/CFDIKG/blob/master/scripts/owlnets/OWLNETS-UMLS-GRAPH-12.py) to convert them into the format that the eleven bases UBKG CSVs are in. Once these CSVs have been created, we append them to the base CSVs. The OWLNETS script was originally written to ingest any OWL-formatted file seamlessly into the UBKG. However, we have since adapted the script to handle any file that follows the format of the nodes and edges files. The format used by OWLNETs can be found at https://github.com/dbmi-pitt/ubkg/tree/main/user_guide.

#### Step 4: Building the graph with the Neo4j Bulk Import tool

Lastly, once the nodes and edges files for each dataset have been converted to CSV format and appended to the base CSVs, we use Neo4js bulk import tool to build the graph on the Neo4j platform. The build process for the current version of Petagraph takes approximately 20 minutes.

### Graph Statistical Analyses

**Graph statistical analyses** were performed with the Neo4j Graph Data Science (GDS) statistics library (v1.6.1) in the Cypher query language. We applied GDS procedures after excluding certain data sources from analyses to show the relative impact of every data source on graph statistics (**Table 2**). Cypher was used to create a Concept graph. Graph centrality and importance were calculated with the functions gds.degree.stats and gds.pageRank.stats (maxIterations: 20, dampingFactor: 0.85). As a measure of community counts in the Concept-Concept graph, the function **gds.labelPropagation.stats** estimated the number of highly connected communities using the Label Propagation Algorithm (LPA). Finally, we utilized **gds.triangleCount.stats** to quantify the global triangle (3-cliques) count in Petagraph and the listed subgraphs.

**Table 1a:**
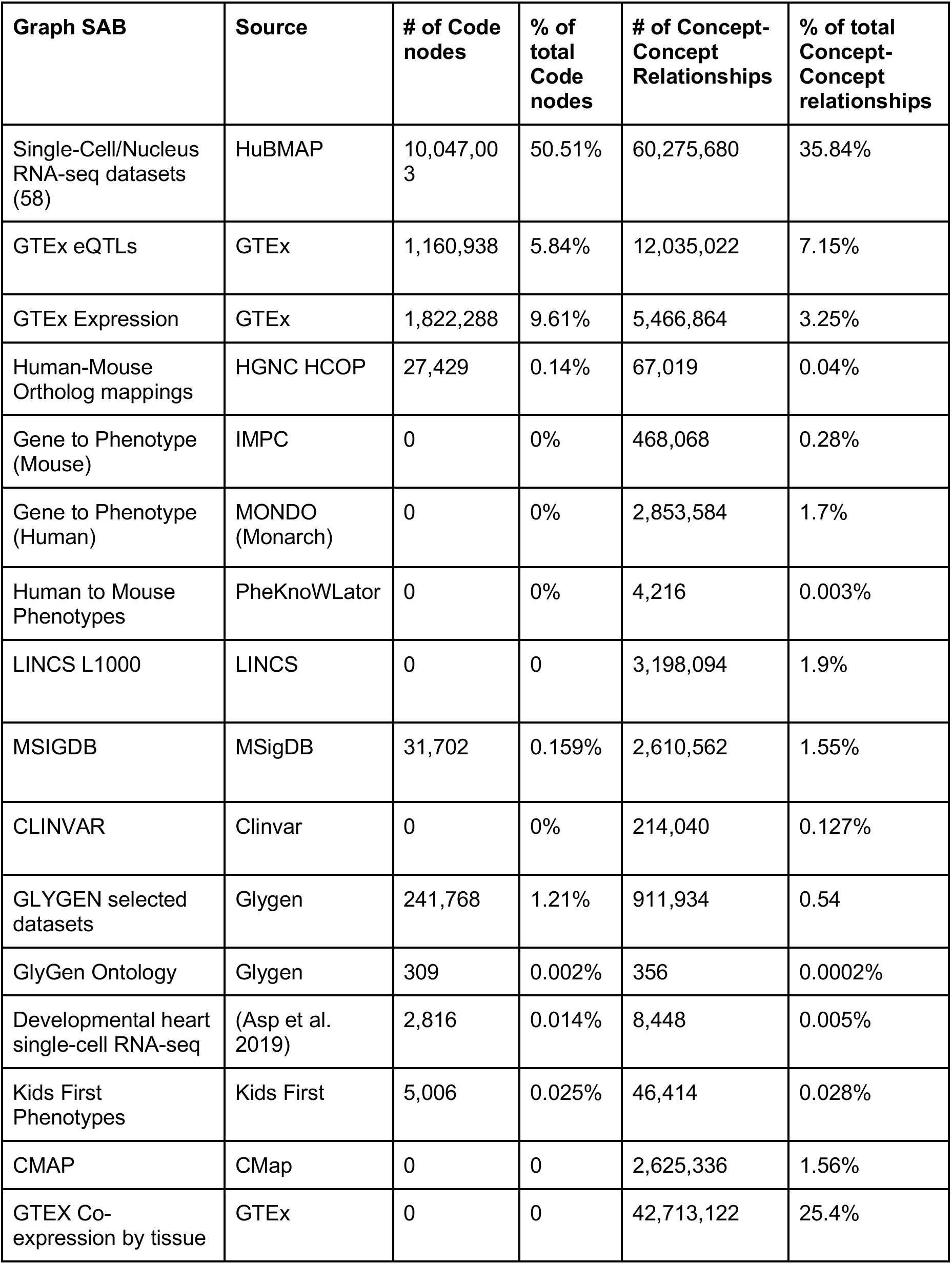
Quantitative datasets and ontologies in Petagraph. * *explain why we’re only reporting Concept (CUI) counts and Concept (CUI-CUI) edges

**Table 1b.** Biomolecular-derived datasets in Petagraph

| **Dataset (citation)** | **Source** | **Graph SAB** | **Has edges with** |
| --- | --- | --- | --- |
| Single-Cell/Nucleus RNA-seq datasets (58) | HuBMAP | HUBMAPSC | HGNC, UBERON, self |
| GTEX eQTLs | GTEX | GTEX_EQTL | HGNC,UBERON,dbSNP, chr loc, P-value nodes |
| GTEX Expression | GTEX | GTEX_EXP | HGNC, UBERON, TPM nodes |
| Human-Mouse Ortholog mappings | HCOP HGNC | HGNC_HCOP | mouse gene nodes, HGNC |
| Mouse Gene-Phenotype mappings | IMPC | IMPC | mouse gene nodes and MP |
| Human Gene-Phenotype mappings | MONDO (Monarch) | MONDO | HGNC, HPO |
| Human-Mouse Phenotype mappings | PheKnowLator | HGNC_HPO | HPO, MP, self |
| LINCS L1000 | LINCS | LINCS_L1000 | ChEBI, HGNC |
| MSigDB | MSigDB | MSIGDB | HGNC |
| ClinVar | Clinvar | CLINVAR | HPO, OMIM, EFO, MONDO, HGNC |
| Glygen selected datasets | Glygen | GLYGEN | Glygen Saccharide, Glycosylation sites, UniProtKB, HGNC |
| Developmental heart single-cell RNA-seq | Asp et al. 2019 | SCHeart_PMID_31835037 | UBERON, HGNC, Expression |
| Kids First Phenotypes | Kids First DRC | KF | HPO |
| Connectivity Map | CMap | CMAP | ChEBI, HGNC |
| GTEX Co-expression by tissue | GTEX | GTEX_COEXP | HGNC |

**Table 2.** : Graph statistics. . Each “Minus” column indicates the graph statistics calculated minus that datasource.

| **Graph Version** | **All CUI-CUI** | **Minus MSIGDB** | **Minus CLINVAR** | **Minus HUBMABSC** | **Minus LINCS** | **Minus GTEX_COE XP** | **Minus GLYGEN** | **Minus CMAP** | **Minus GTEXEQTL** |
| --- | --- | --- | --- | --- | --- | --- | --- | --- | --- |
| **Relationship Count (difference vs full graph)** | 168155890 | 165545328 (2610562) | 167941850 (214040) | 107880210 (60275680) | 164957796 (3198094) | 125442768 (42713122) | 167243956 (911934) | 165530554 (2625336) | 156120868 (12035022) |
| **Global Triangle Count** | 7503683363 | 7495671978 | 7503038504 | 7503683356 | 7467533393 | 13440416 | 7503682261 | 7483258009 | 7502319751 |
| **Label Propagation Community Count** | 213083 | 213366 | 212830 | 212798 | 211410 | 214850 | 214011 | 212551 | 2178933 |
| **Mean Centrality Score (Page-Rank)** | 0.958 | 0.958 | 0.958 | 0.529 | 0.958 | 0.958 | 0.948 | 0.958 | 0.874 |
| **Max Centrality Score (Page-Rank)** | 208847 | 211539 | 208941 | 69823 | 211340 | 228403 | 208852 | 211484 | 207328 |
| **Mean Degree Centrality** | 8.850 | 8.713 | 8.839 | 5.678 | 8.682 | 6.602 | 8.802 | 8.712 | 8.213 |
| **Max Degree Centrality** | 1686840 | 1686840 | 1686840 | 737252 | 1686840 | 1686840 | 1686840 | 1686840 | 1686840 |

**Semantic type heatmaps** were generated from a selection of 54 (out of 127) Semantic Types in Petagraph, where the selected types are related to anatomy, phenotypes, diseases, and chemical species. The distinct relationships in each Semantic Type heatmap (**Figure 5**) were also retrieved from the graph. For **Figures 5b and 5c**, the frequency of such relationships are shown between pairwise combinations of semantic Types, connected through their Concept nodes. Therefore Concept-Concept relationships where one or both Concept nodes are not connected to Semantic Types are excluded. The presence of relationships (**Figure 5a**), the cumulative number of relationship types (**Figure 5b**), and the relationship counts (**Figure 5c**) were plotted using the package *pheatmap* v1.0.12 (Kolde 2019) in RStudio v 1.4.1106 (Posit team 2022) R v4.0.4 (R Core Team 2022). To increase the dynamic range in Figures 5b and 5c, the base10 logarithm was used, and to avoid log_10_ (0), we added 1 to all values.

**Figure 3:**
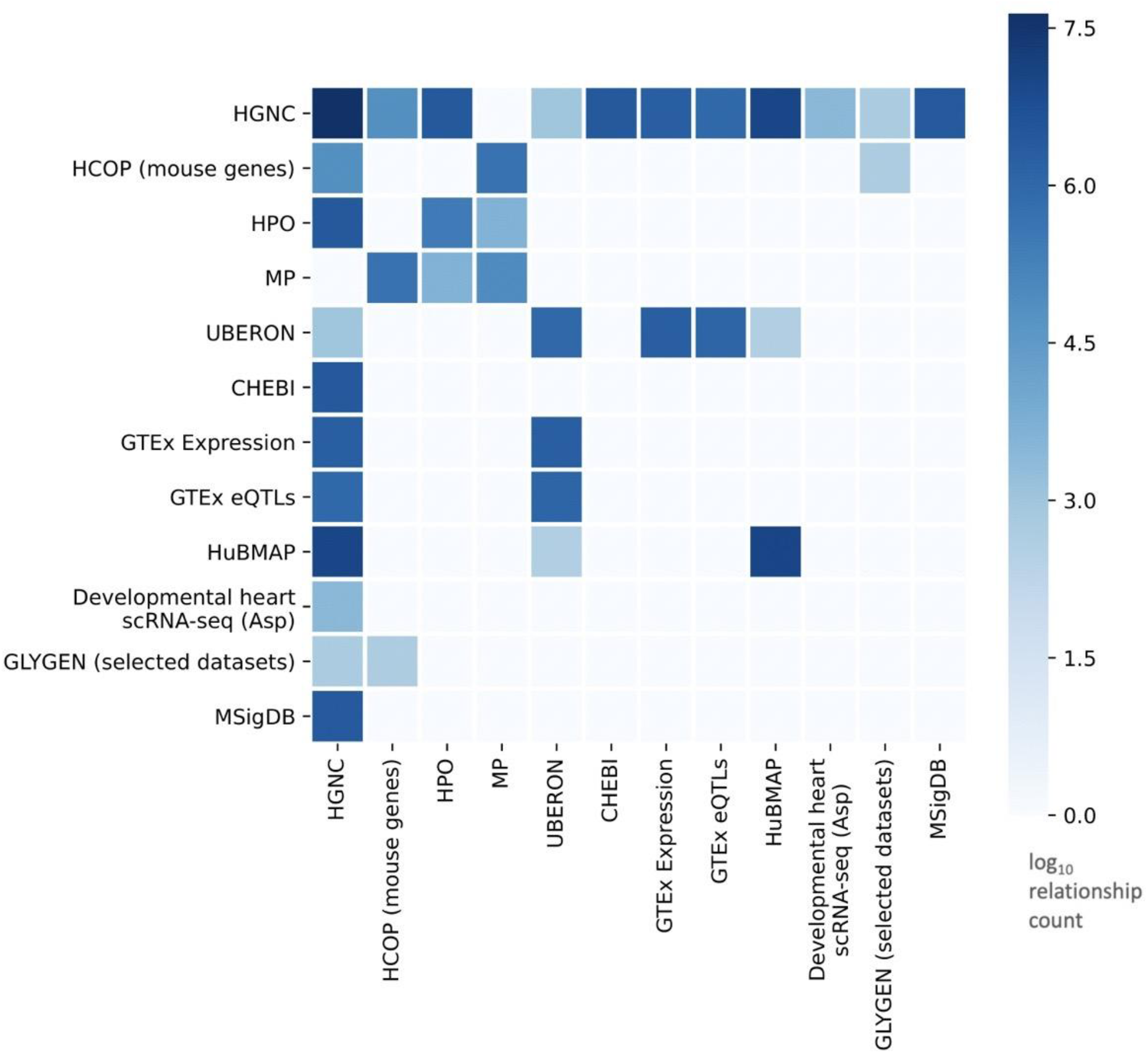
Dataset interconnectedness. Relationship counts between selected data sources in Petagraph. Several ontologies that are part of the UBKG ingestion are shown (HGNC,UBERON,HPO, CHEBI). The integrated datasets are largely gene-centric with most datasets having relationships to HGNC nodes

**Figure 4:**
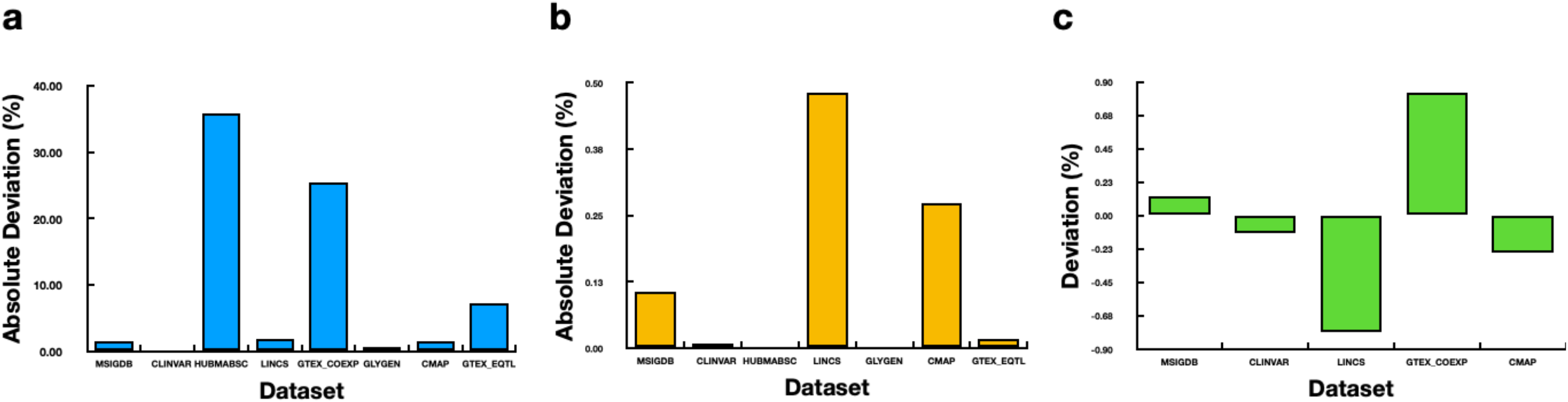
Graph Statistics by ingested dataset. Absolute deviations from the mean for selected datasets showing: **a)** relationship counts, **b)** global triangle counts and **c)** label propagation community count.

**Figure 5:**
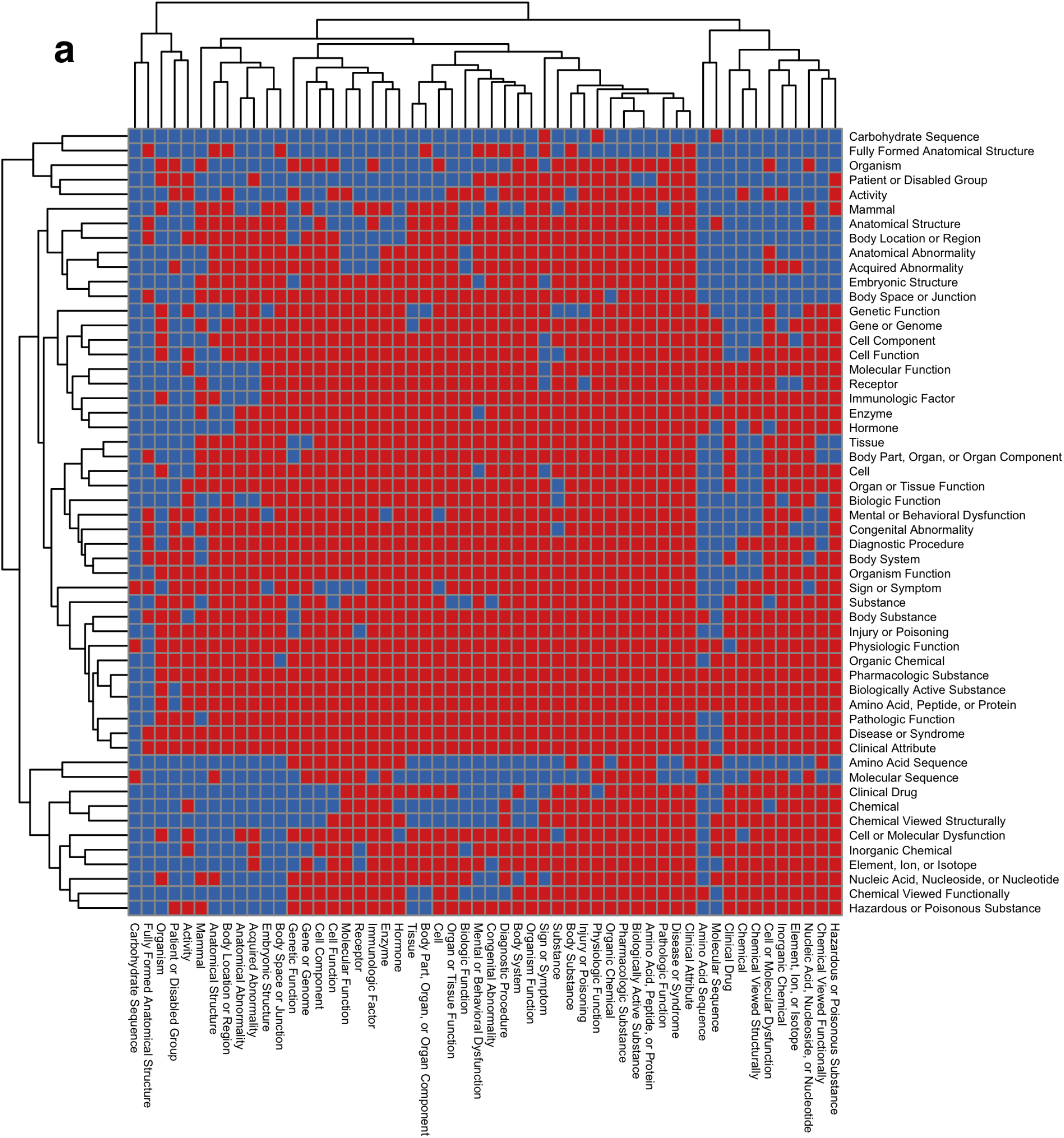
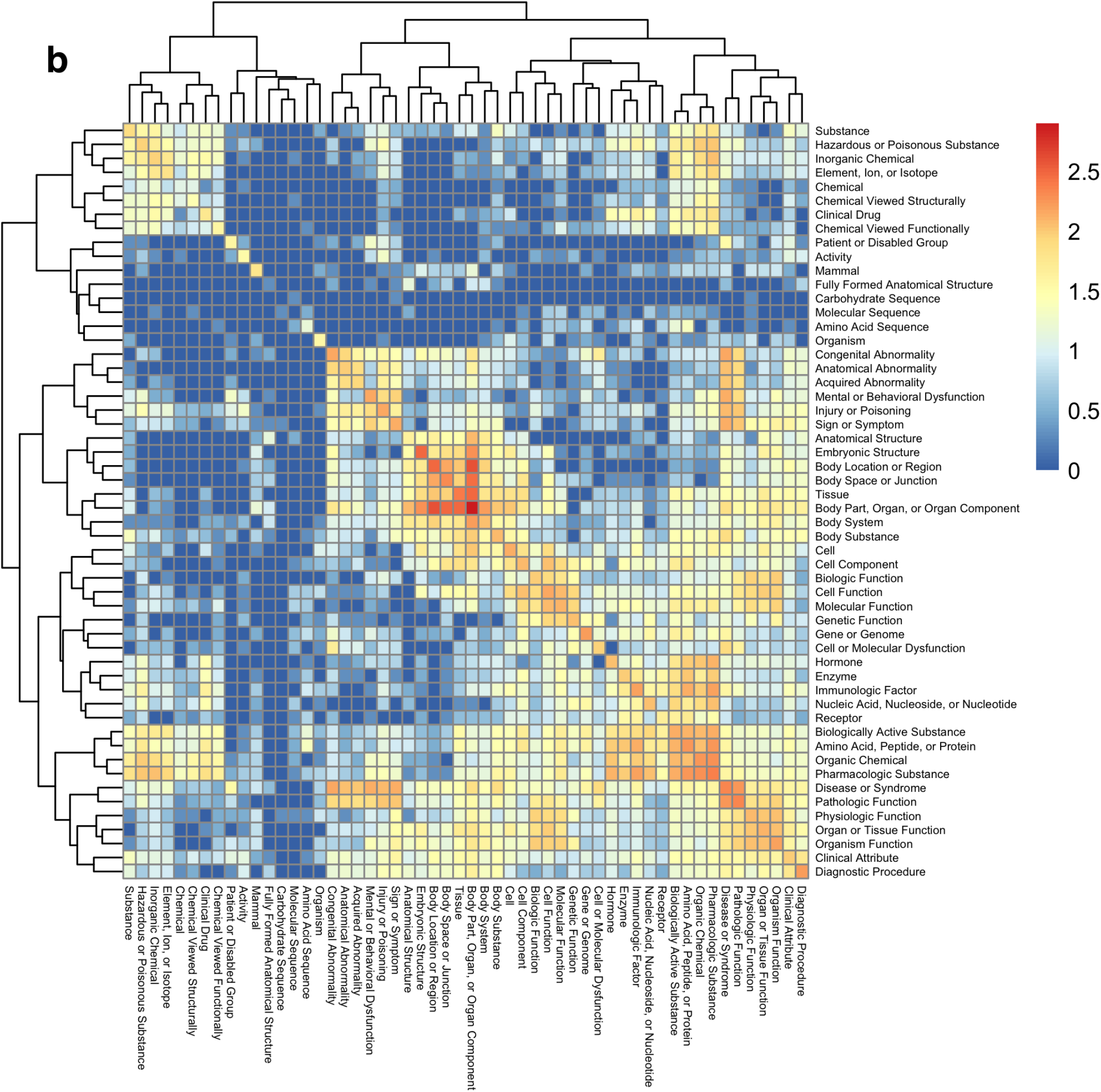
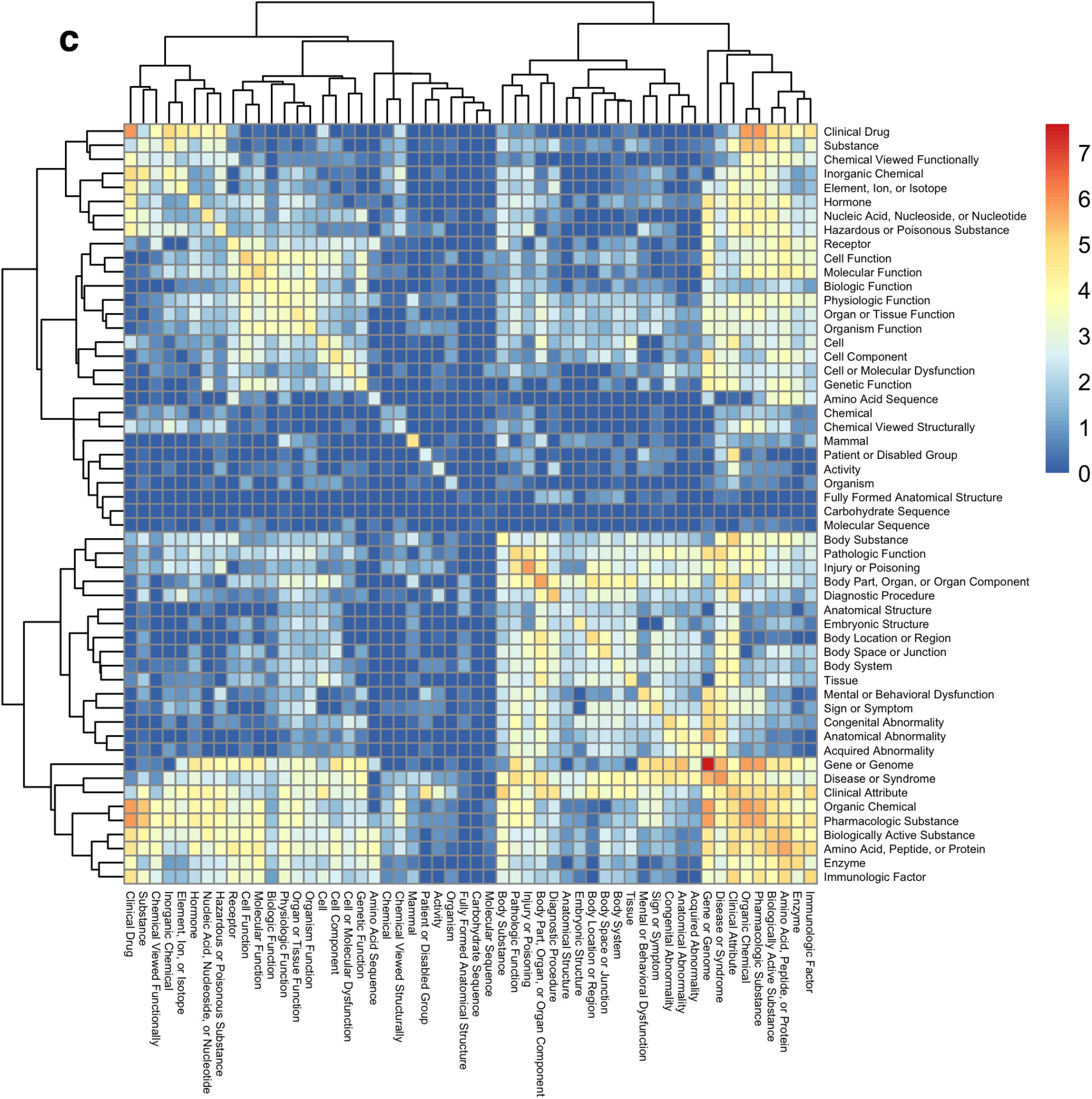
Heatmap representation of relationship statistics of 54 selected. Semantic Types. In all graphs, data represents Concept-Concept node connections. **a)** the presence (red) or absence (blue) map of at least one relationship (type+SAB) between pairwise combinations of Semantic Types shows. **b)** The color intensity representation of the diversity of the log10 relationship counts (type+SAB) connecting the candidate Semantic Types. **c)** The color intensity representation of the log10 number of outgoing relationships (from row to column) connecting the candidate Semantic Types. Note that for **(a)-(c)** the matrices are diagonally symmetrical as Concept-Concept relationships are bidirectional.

#### Shortest Path Length Analysis

Pairwise shortest path lengths between 256 million distinct pairs of **`HGNC`** concept nodes were calculated using a Petagraph subgraph consisting of only Concept nodes and excluding **`HUBMAPSC`** data. Analysis was performed using the distances function from *igraph* v1.2.11 (Csardi et al. 2006) in RStudio 2022.12.0.353 (Posit team 2022) and R v4.2.2 (R Core Team 2022). **Figure 6** shows that the majority of shortest paths fall within the range of 1 to 10.

**Figure 6:**
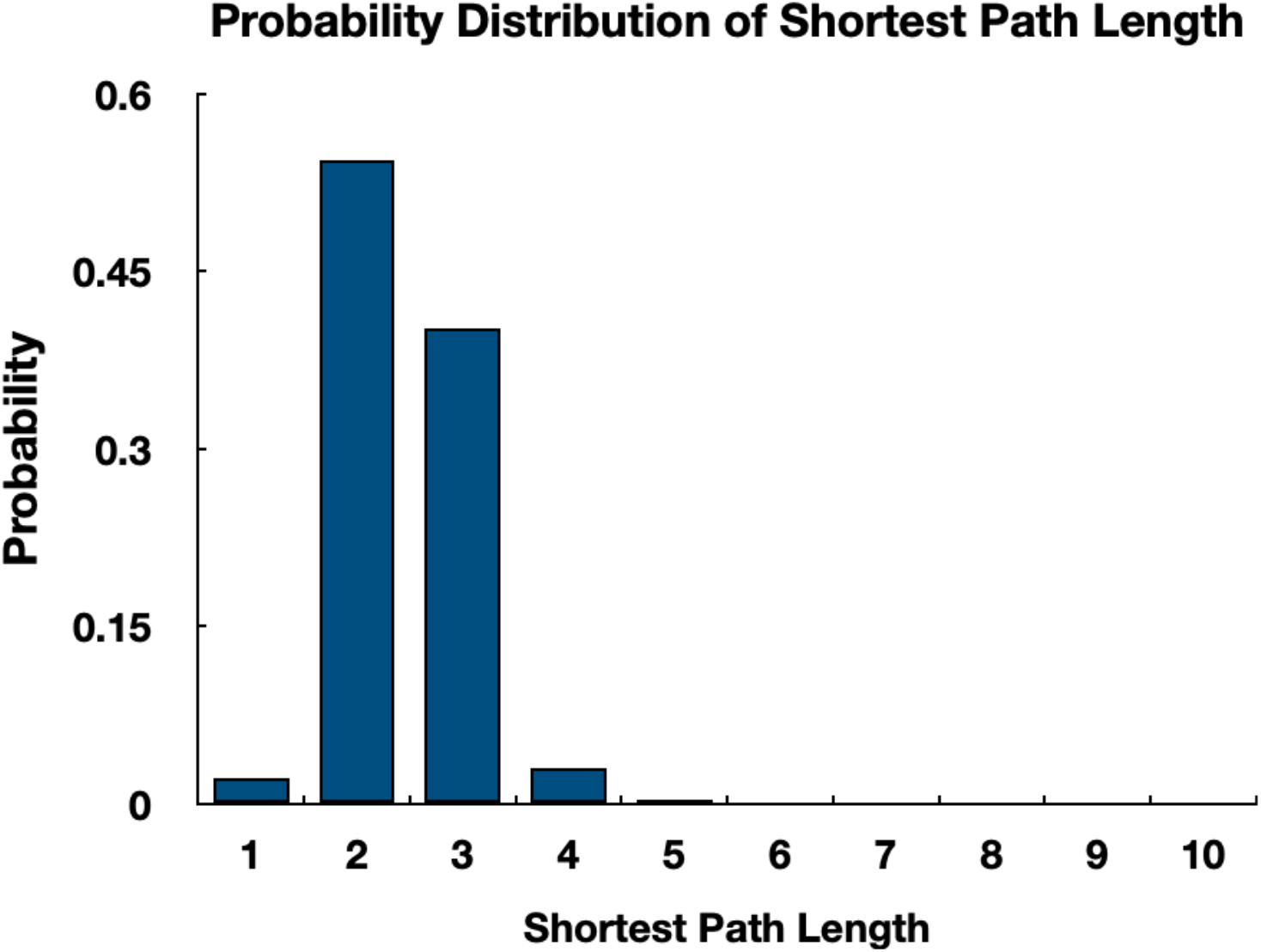
Shortest Path Lengths. Probability distribution of associated Concept to Concept node shortest path lengths for 256 million pairwise combinations of human gene Concept nodes in Petagraph.

### Use Cases

#### eQTL vs phenotype Use Case

GTEx eQTLs represent variants that are “expression Quantitative Trait Loci” (eQTL) that can potentially indicate variants that have upstream regulatory influences over a downstream target gene. We sought to identify a possible link between animal model phenotypes and eQTLs by linking a query across the graph. Figure 7b shows the neo4j Cypher query to generate all eQTLs associated with a particular mouse phenotype, for example mammalian atrial septal defect (MP:0010403). The query matches all child phenotypes below the level of MP:0010403 using a recursive search on the graph and extracts all genes associated with those phenotypes. Gene-to-phenotype links from mice were included from IMPC using the OBO Relations Ontology (RO) RO:0002331 (involved in) term (OBO_Foundry). The mouse gene-human gene relationships were included in the graph from the **`HGNC HCOP`** resource <u>(Yates et al. 2021)</u>. The links between genes and eQTLs and the p-values for each eQTL-tissue relationship were provided by GTEx. In the query we selected eQTLs for only the heart left ventricle myocardium using the Uberon ontology (UBERON: 0006566) and right atrium (UBERON: 0006631) as these anatomical parts are associated with atrial septal defects <u>(Mungall et al. 2012)</u>. The result of the query yielded 309 human eQTLs with p<0.05 that are potentially associated with genes linked to mammalian atrial septal defects (MP:0010403).

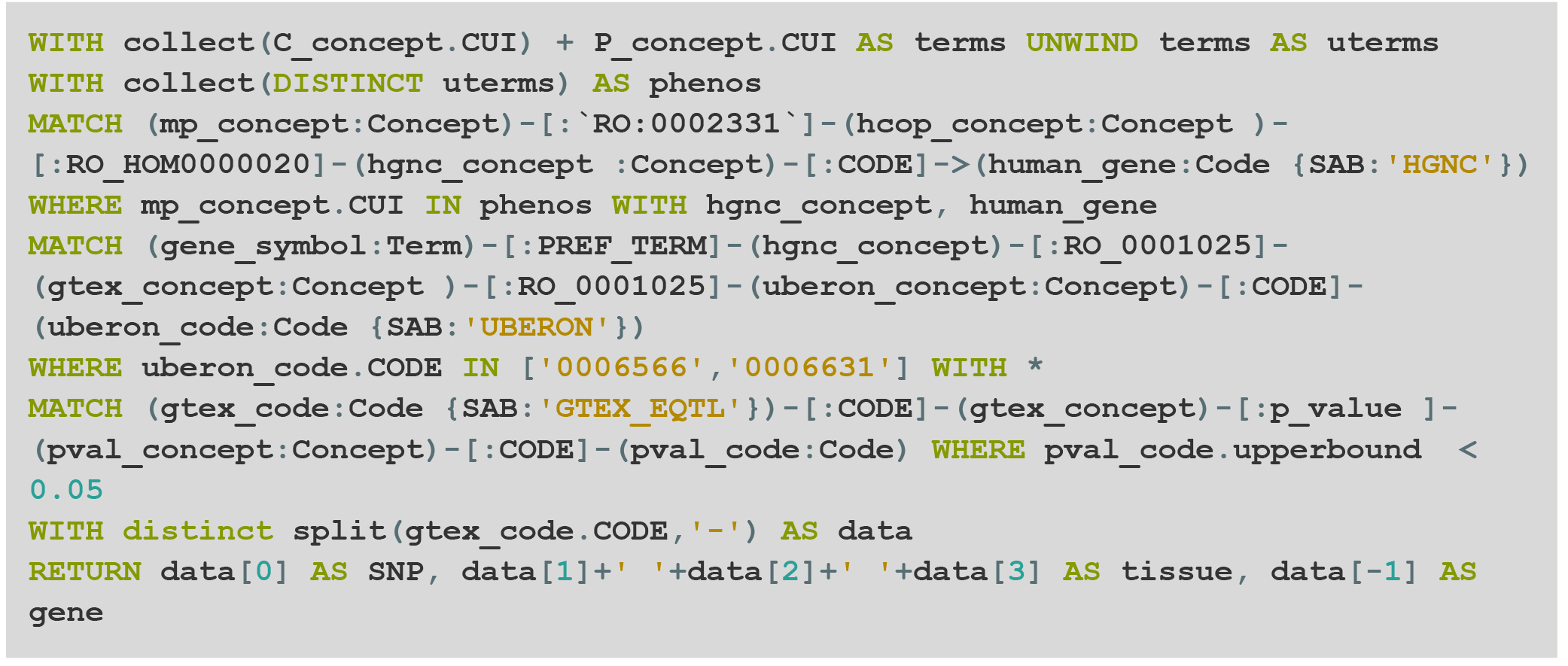

**Figure 7:**
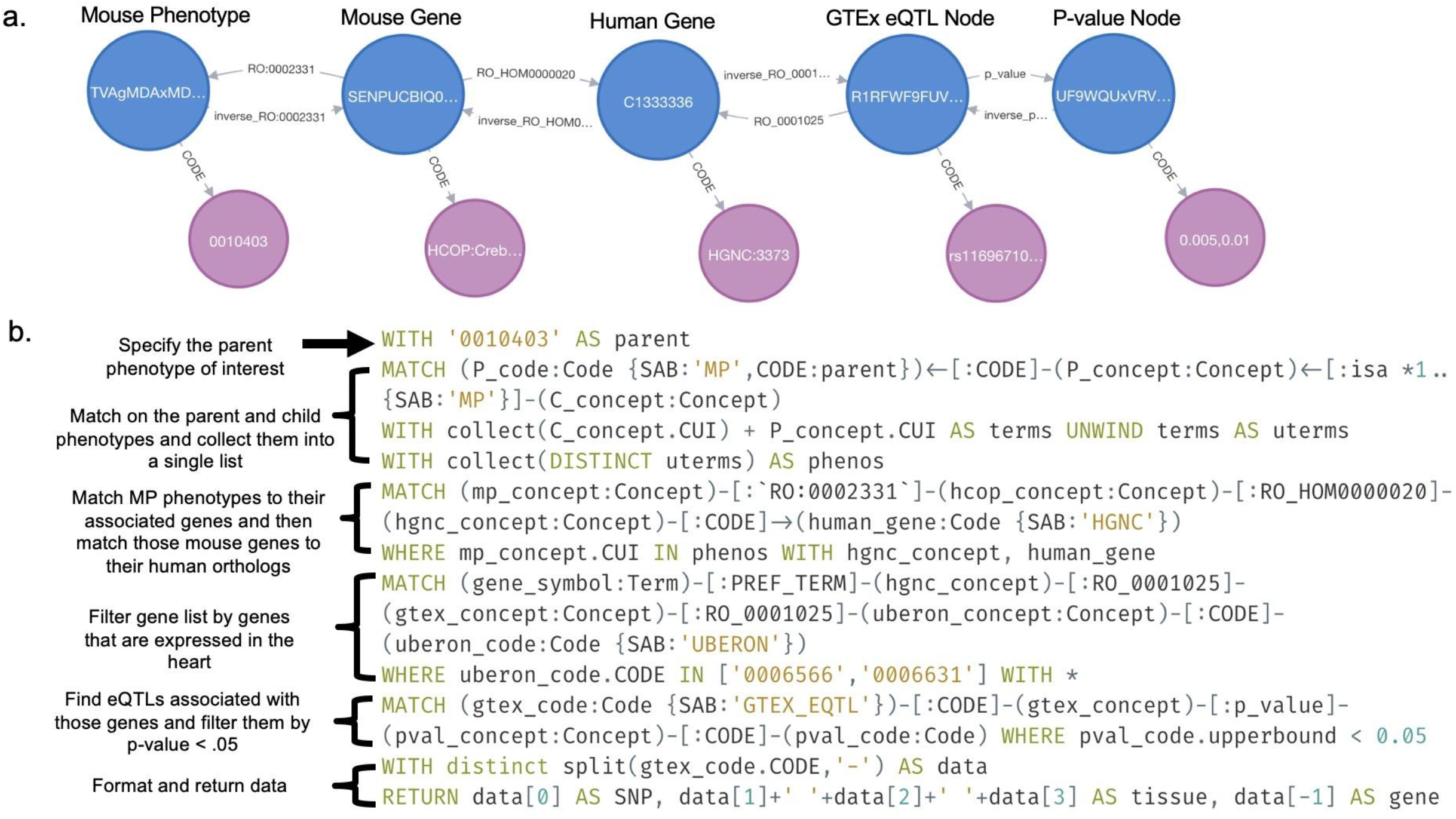
Recursive queries on ontologies allow for returning results and increasing search depths. This is an example of a recursive (transitive closure) search using the mammalian phenotype (MP) **Atrial Septal Defects** as the parent phenotype. This query returns all of the eQTLs (p-value < .05) from ASD (**`MP:00110403`**) as well as its child phenotypes, based on genes that are expressed in the GTEx heart data. Concept nodes are in blue, code nodes in purple, and term nodes in green.

#### Glycosyltransferase evaluation Use Case

The GlyGen project provides resources for studying biology related to glycosylation (York et al. 2020). GlyGen’s human glycosyltransferase proteins dataset was utilized to obtain median TPM expression levels for tissues in the GTEx expression dataset. The expression levels (upper bound and lower bound) for the genes were returned from the graph. In total, 220 glycosyltransferases were plotted for 40 individual tissues. The GlyGen dataset provided a list of glycosyltransferases which were integrated into the graph by simply adding a **`is_glycosyltransferase`** relationship between a (human) glycosyltransferase Concept node and the corresponding HGNC Concept nodes. Then the relationship **`RO:0002206`** (expressed_in) between HGNC and GTEx Expression Concept nodes was utilized to connect to the GTEx Expression nodes. Lastly, Expression Bin nodes, which bin GTEx Expression median TPM values between an upper and lower bound are accessed through a **`has_expression`** relationship. All TPM values were returned in this example, however it would be trivial to add to the query to filter TPM values for a certain cutoff, or return only the maximum TPM and the associated glycosyltransferase. The returned data can then be downloaded directly from the Neo4j browser or programmatic access to the graph can be established using Neo4j’s drivers and the data can be returned directly into your environment. Python 3.7 and matplotlib 3.4 were then used to create Figure 8.

**Figure 8:**
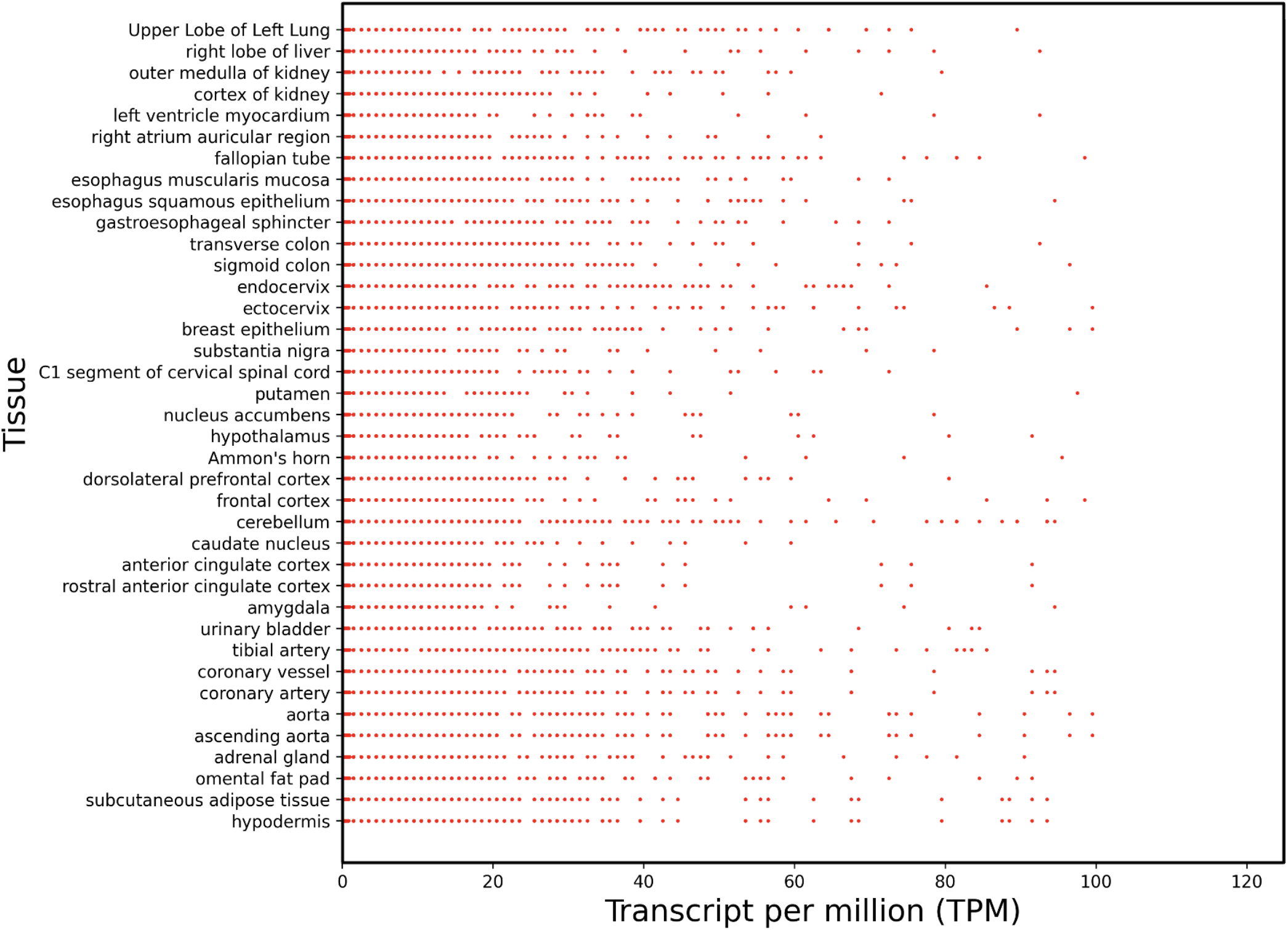
Expression levels for the human glycosyltransferase gene family. Each dot represents a glycosyltransferase gene’s TPM within each GTEx tissue. Values are medians for that gene calculated within each tissue cohort.

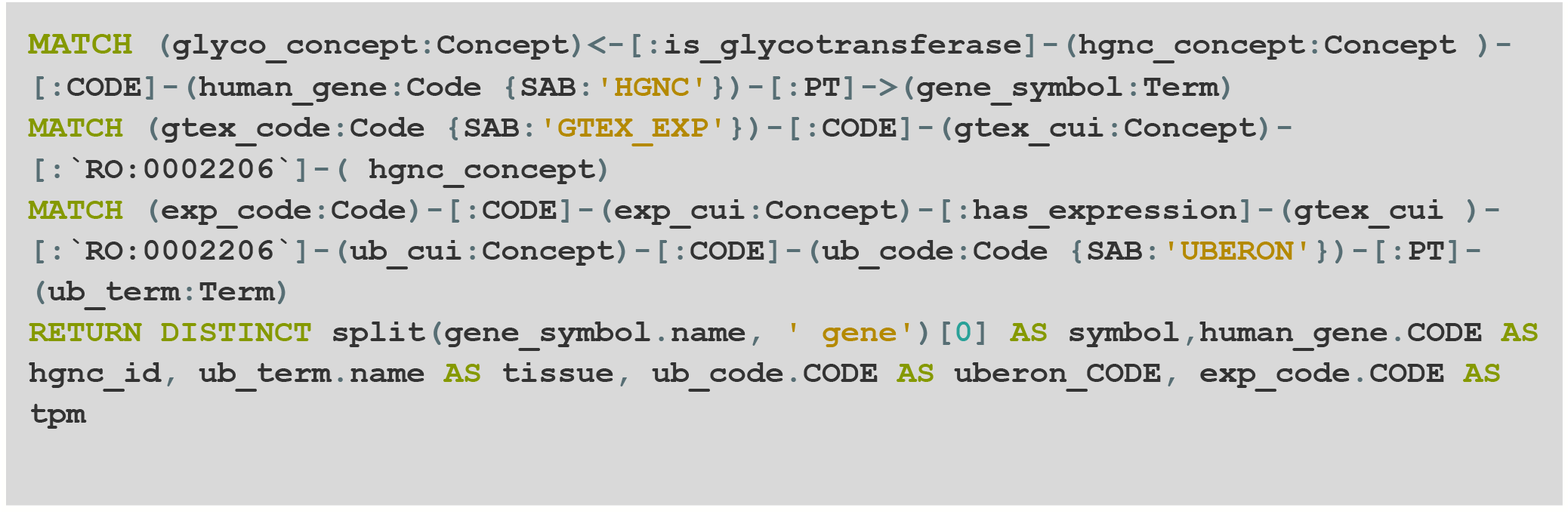

#### Glycosylation by Phenotype Use Case

We obtained a set of glycosylated proteins, glycosylation types (O-linked, N-linked) and protein glycosylation sites associated with common human congenital heart defects (CHDs) through querying the knowledge graph using two different methods, one for human using the Human Phenotype Ontology (HP) (Köhler et al. 2021) and mouse using the Mammalian Phenotype Ontology (MP) (Smith and Eppig 2009).

##### Glycosylation by Human Phenotypes (HPO)

To identify glycosylated proteins and tissues in humans relative to disease status, we queried Petagraph with Concepts in HP with the following Codes: HP:0001629 (Ventricular Septal Defect, VSD), HP:0001631 (Atrial septal defect, ASD), HP:0001643 (Patent ductus arteriosus, PDA), HP:0001636 (Tetralogy of Fallot, ToF), HP:0001680 (Coarctation of the aorta, CoA), HP:0001719 (Double outlet right or left ventricle, DORV), HP:0001650 (Aortic valve stenosis, AVS), HP:0012303 (Aortic Arch Anomaly, AAA), HP:0006695 (Atrioventricular Septal Defect AVCD), HP:0001660 (Truncus arteriosus, TA), HP:0030853 (Heterotaxy, HTX), HP:0001718 (Mitral valve stenosis, MS). For each HP Concept, all “child nodes” below the originally selected HP phenotype were identified down to the terminal branches. All resulting HP terms were associated with genes identified on the HP website which publishes relationships of phenotype to gene features in their `phenotype_associated_with_gene` table. This and the `genes_associated_with_phenotype` tables derive gene-phenotype relationships by linking genes in `OMIM` and estimated phenotypes to disease, as discussed (Peter N Robinson, Sebastian Köhler, Sandra Doelken, Sebastian Bauer 2022). We use relationships connecting data from UniprotKB (`UNIPROTKB`) Concepts called `has_product` that links gene names (HGNC Concepts) to `UNIPROTKB` protein IDs, respectively <u>(Boutet et al. 2016)</u>. We then used the relationships `on_protein`, and `has_glycan` to identify the glycosylation sites on each protein. Therefore, using this relationship set, we were able to link across phenotypes, genes, proteins, and glycosylation sites, and identified or predicted glycans (saccharide) Concept nodes.

##### Glycosylation by Mouse Phenotypes (MP)

Mouse phenotype-to-gene information was identified from IMPC and MGI. The resource links phenotypes to genes based on observed gene perturbations. Mouse phenotypes as MP identifiers from these resources were identified as equivalent to HP terms of human phenotypes of interest: i.e. MP:0010402 (Ventricular Septal Defect, VSD), MP:0000284 (Double outlet right ventricle, DORV), MP:0004113 (Aortic Arch Anomaly, AAA), MP:0010454 (Truncus arteriosus, TA), MP:0010403 (Atrial septal defect, ASD), dMP:0010412 (Atrioventricular Septal Defect, AVCD), MP:0004133 (Heterotaxy, HTX), MP:0004110 (Dextro-Transposition of the Great Arteries, TGA), MP:0010449 (Right ventricular outflow tract obstruction, RVO), MP:0010413 (Complete atrioventricular canal, CAVC), MP:0003387 (Coarctation of the aorta, CoA).

After the mouse gene list was obtained, the human orthologs of the extracted mouse genes (`has_human_ortholog`) were linked using the `HGNC HCOP` relation (Yates et al. 2021) and then retrieved the glycans and glycosylation type/sites in the proteins encoded by those genes associated with the phenotypes above, following the steps described in the last paragraph.

Once the data was extracted, we computed the ratio of genes with glycosylated proteins to all genes associated with each phenotype to estimate how much the involved gene product would interact with glycosylation in general. Additionally, we found the relative number of distinct glycans per glycosylated proteins as a measure of the diversity of saccharides employed to modify such proteins. Finally, we took the average number of glycosylated types/sites per glycosylated proteins for each phenotype as well as all human genes to measure the extent of glycosylation in each phenotype group. The results of these analyses are reflected in **Figure 9**.

**Figure 9:**
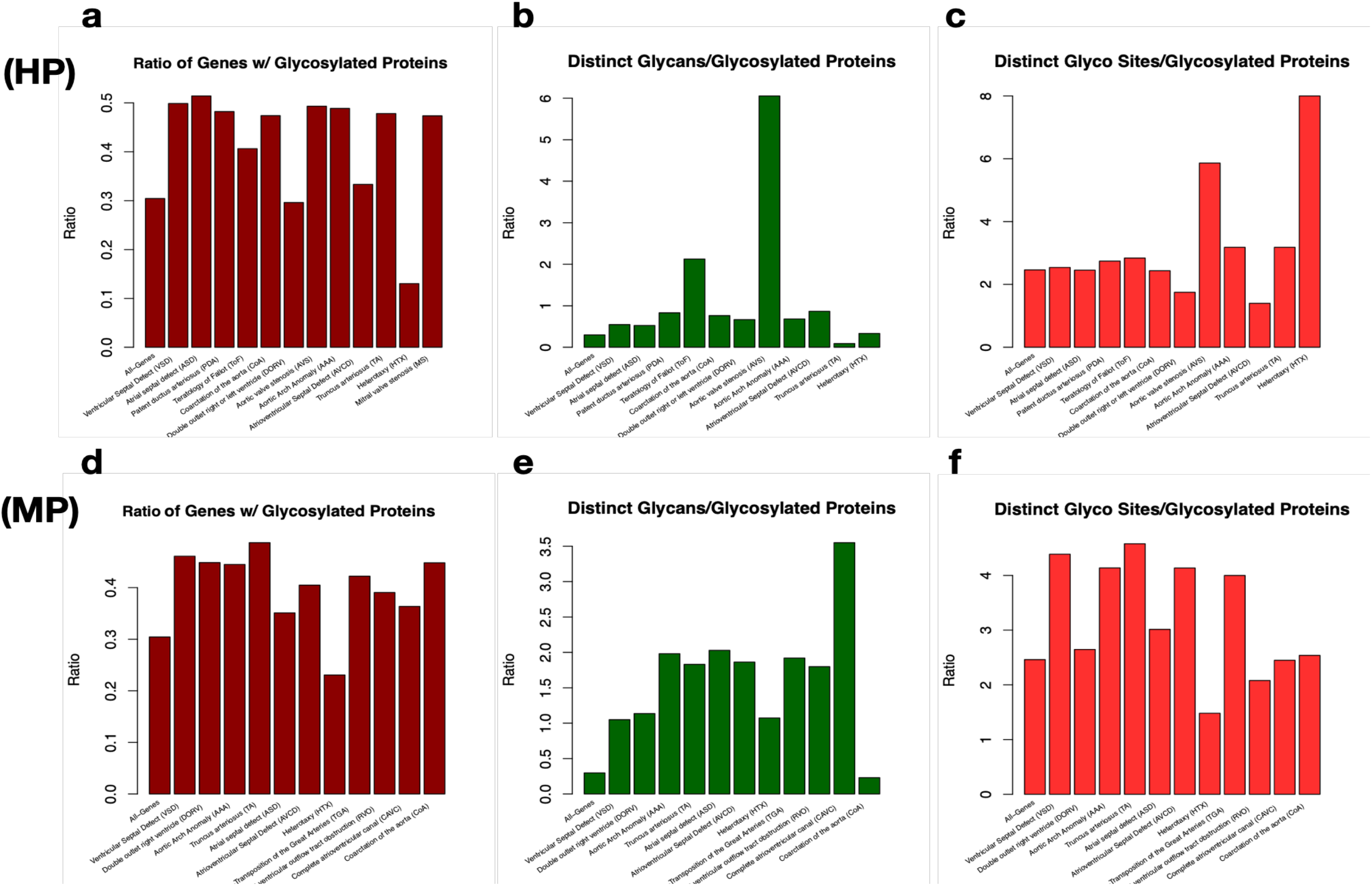
Query results on heart-defect phenotype-associated glycosylation targets. The query linked phenotypes from from **a-c)** OMIM (human) and **d-f)** IMPC (mouse) to associated genes, proteins, glycosylation sites and glycans using the intersection of UniCarb-DB, GlyConnect, and O-GlcNAc (see Methods). The bars represent the ratios of known glycosylated genes vs all source genes associated with each condition, showing the relative depletion of glycosylated sites by type of heart defect. Heterotaxy, for example, is much lower in glycosylated gene products associated with the disorder (**a** and **d**). CHD-associated genes have higher diversity of glycans/proteins (**b** and **e**) and higher sites/proteins (**c** and **f**).

#### Developmental origins of heart defects use case

We utilized previously published single-cell gene expression data from embryonic heart samples at 6.5-7 post-conception weeks (PCW) by gene ID, cell (cluster) identity, anatomy, and cluster p-value and fold change enrichment values (Asp et al. 2019). We used the Cell Ontology (Osumi-Sutherland et al. 2021) to link cell type to cluster identity. For those cell types not found in the Cell Ontology, we created new ‘temporary’ types within Petagraph until such time that the Cell Ontology is updated. There were 14 author-defined cell types (clusters). We used the Seurat package in R to further analyze the single-cell RNA-seq data with differential expression analysis (Butler et al. 2018; Stuart et al. 2019; Hao et al. 2021). By clustering the developing heart cells by the author-defined cell types we could identify differentially expressed gene (DEG) markers that provided p-value and log2-fold-change values representing the enrichment of a DEG marker in a cluster (cell type) versus all other clusters. Nearly 3,000 biomarkers were identified with p<0.05 and log_2_FC>1 across the 14 cell types. These biomarker data were mapped to human gene CUIs by HGNC ID and the DEG biomarkers were created as new Concept nodes that are uniquely identified by their relationships to the biomarker (gene) and cell type (CL) Concept nodes. We then link the single-cell data in developing the human heart to the mouse phenotypes of developmental heart defects through MP.

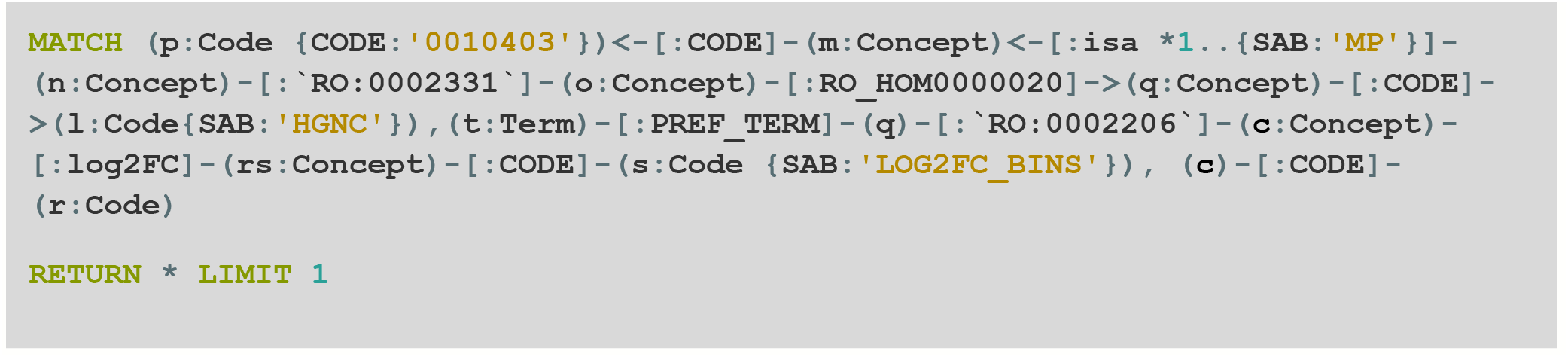

#### Rofecoxib use case

For this use case, we seek to link an expression signature for the drug rofecoxib (**`CHEBI:8887`**) to different GTEx tissue expression profiles. The gene expression tissue profiles for CMap and LINCS L1000 data were summarized for use in Petagraph as discussed in the Supplemental data in the “LINCS L1000” section. The summarized gene expression profile for rofecoxib from CMap and L1000 (**`CMAP`** or **`LINCS L1000`**) (Subramanian et al. 2017) was compared to the tissue-specific gene expression profiles from GTEx. The ratio was calculated for genes in rofecoxib profiles found in GTEx tissues (in **`GTEX_EXP)`** with the total number of genes in the tissue with TPM>5. The ratios for each tissue were normalized to the tissue with the highest value. The tissues were then ranked accordingly (Figure 11).

**Figure 10:**
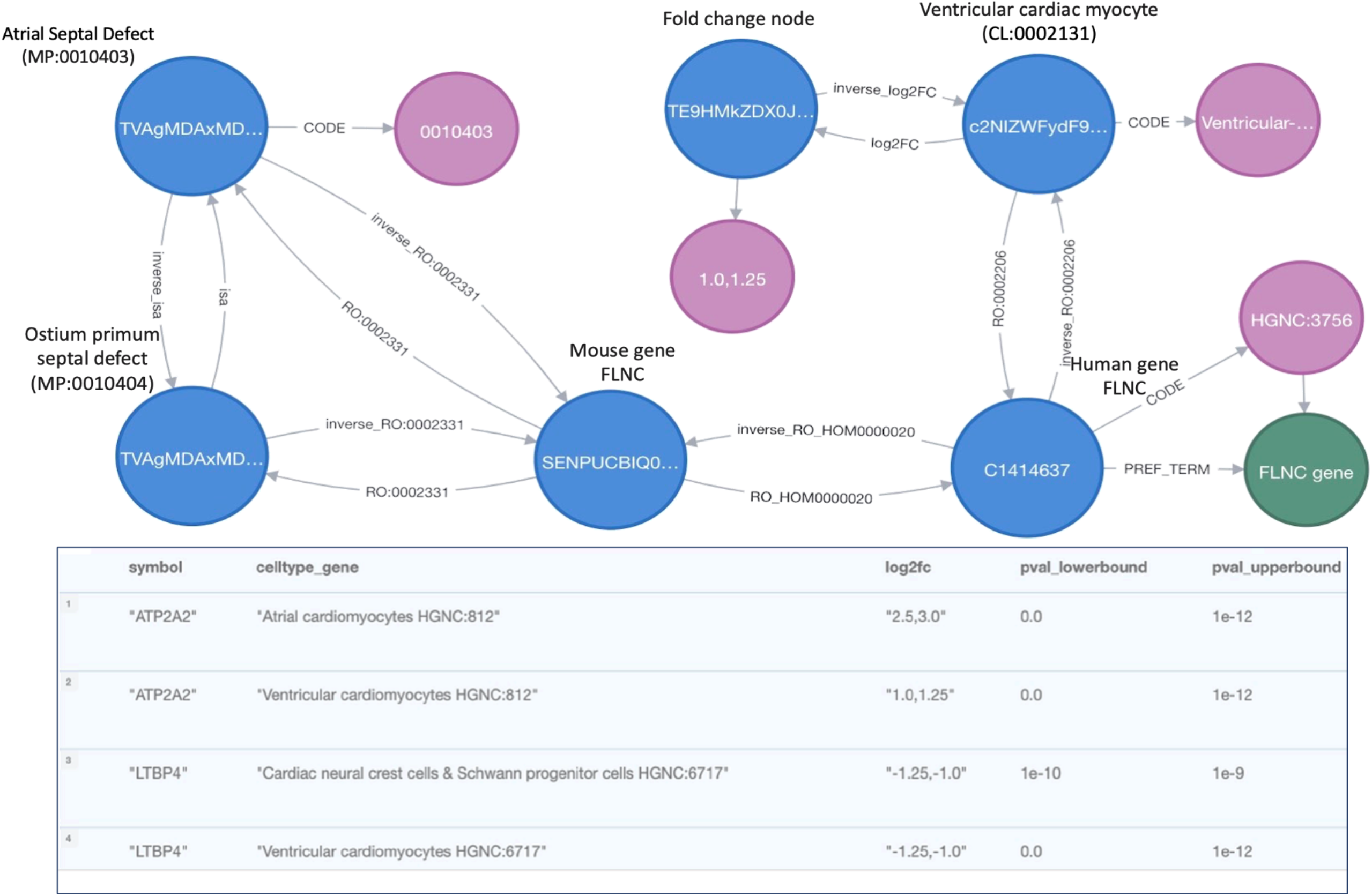
Result of a query linking genes associated with a phenotype to specific cell types in a single-cell 6w human embryonic heart dataset (Asp et al. 2019). The top panel shows a single result of the query, where the gene FLNC, associated with a phenotype Atrial Septal Defects, MP:0010403 is linked to the ventricular cardiomyocyte cell type. The bottom panel shows tabular query results where specific gene markers associated with MP:0010403 were identified for cell type clusters in the dataset.

**Figure 11:**
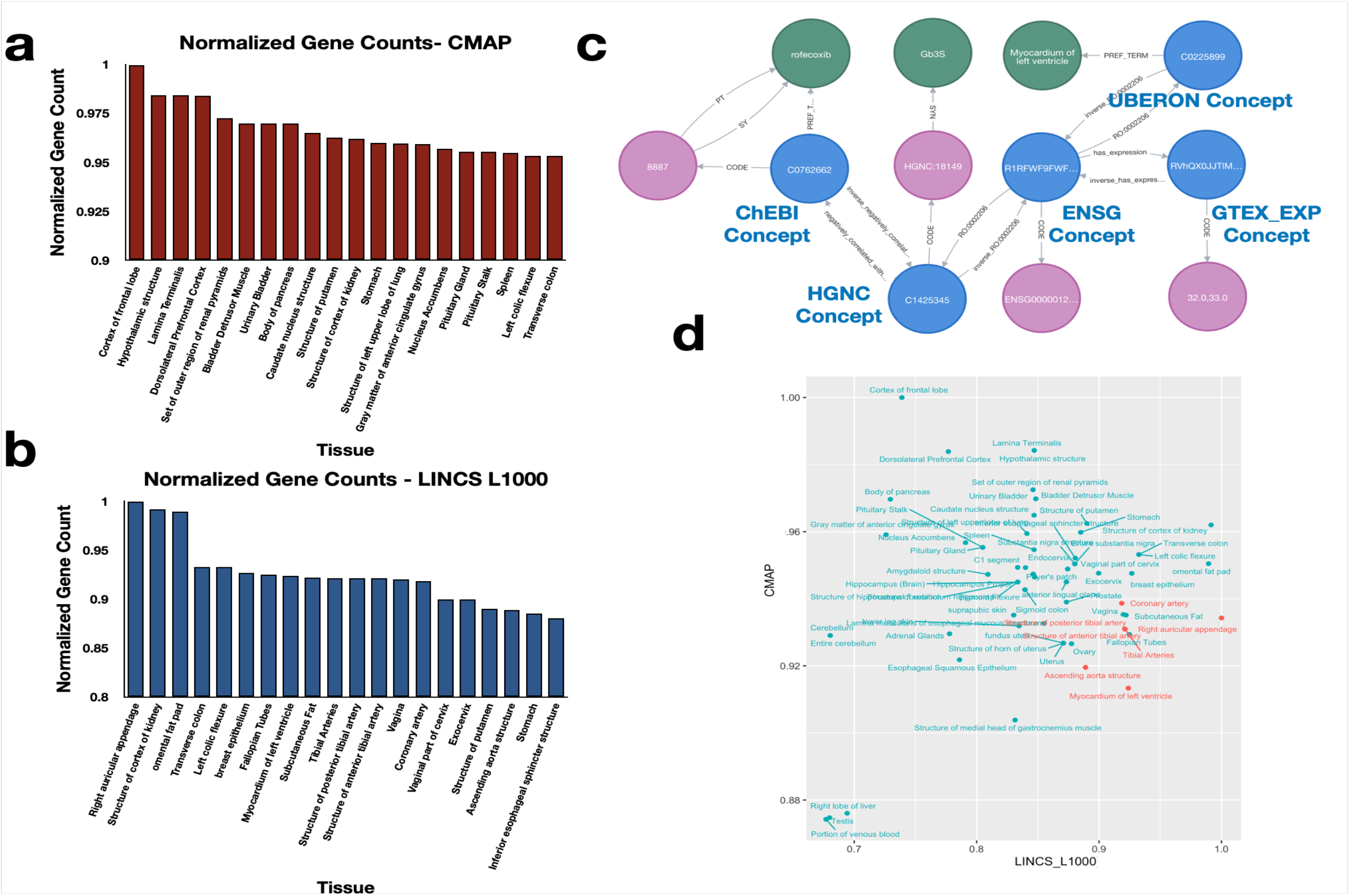
Querying possible drug-tissue interactions for rofecoxib using CMAP, LINCS L1000 and GTEX_EXP datasets. **a)** Top 20 tissues with the highest ratio of genes correlated with rofecoxib (**`CMAP`**) and to all genes (TPM>5), **b)** Top 20 tissues with the highest ratio of genes correlated with rofecoxib (**`LINCS`** L1000) to all genes (TPM>5). **c)** An example query result for possible interaction of rofecoxib and myocardium of left ventricle through Gb3S gene modulation. **d)** Distribution of normalized values of gene interactions per tissue correlated with rofecoxib based on CMAP vs. LINCS L1000 data (cardiovascular system noted in red) .

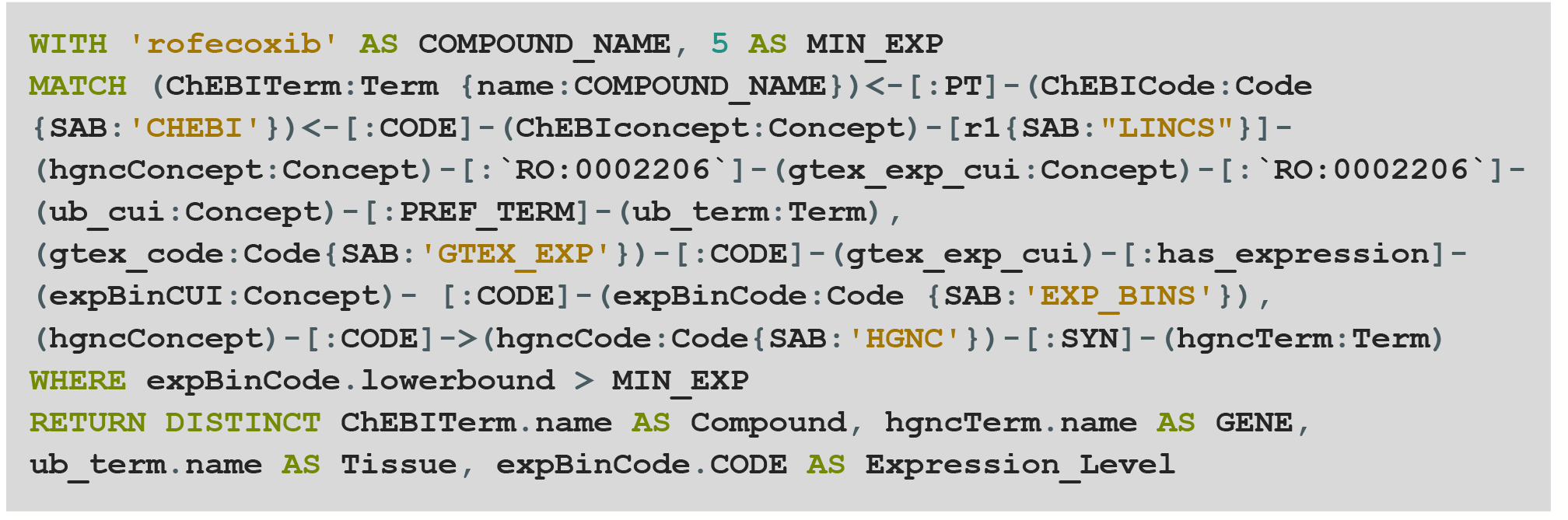

## Results

Data modeling was a major consideration in how we introduced quantitative biomolecular data within the ontology-rich environment supported by the UBKG. Quantitative values are supported within the Petagraph model and can be used and queried as numbers. However, we also wanted to integrate quantitative data within the categorial structure that is native to the UMLS to support complex queries that would not have the overhead for multidimensional, numerical searches. We created Concept nodes representing interval bins of numerical ranges, for example for p-values, expression TPM and log_2_FC. Data points can then be assigned to those bins, which allows for rapid selection of numerical values in the graph.

To demonstrate the contribution of individual datasets to Petagraph’s final structure, we performed a set of graph theoretic statistical analyses of Petagraph and the corresponding subgraphs of Petagraph, each excluding one of the data sources, as shown in **Table 2**. The analysis included an estimation of the relationship counts for each dataset as a subgraph (hence the difference in the number of links between subgraphs), degree centrality analysis, PageRank, global triangle count, and label propagation community count. **Table 2** shows that the majority (>99%) of all triangles in the graph result from the integration of GTEx Coexpression links; that is, 7,490,242,947 triangles within the graph consist of **`HGNC`** Concept nodes with an SAB of **`GTEX_COEXP`**. In understanding this number, one should consider the reciprocal nature of the Concept-Concept relationships as well as the fact that each triangle is counted thrice to account for each vertex. Another notable observation is the contribution of the HuBMAP single-cell dataset, (**`HUBMAPSC`**) to the overall connectivity in the graph, which is reflected in the reduction of the mean degree centrality in the subgraph lacking edges from **`HUBMAPSC`**, which is mainly due to its large number of edges. There are differential numbers of relationships between the datasets. Preferential relationships between different datasets are not explicitly shown here, including several relationship-only data sets. For example, the **`GTEX_COEXP`** data makes up the majority of **`HGNC`**-to-**`HGNC`** relationships and the **`CHEBI`**-to-**`HGNC`** relationships are made up of **`LINCS`** and **`CMAP`** data.

For the fifteen new datasets ingested into Petagraph, we were interested in how those datasets are inter-related by dataset identity. **Figure 3** shows the log_10_ count of relationships across the datasets. The HGNC (human gene) category is directly connected to each human biomolecular-derived dataset. HuBMAP currently has the most human gene relationships given Petagraph’s current model. Future releases of Petagraph will reduce the number of HuBMAP data points through more aggressive summarization. The GlyGen dataset includes glycosylated information on mouse proteins and maps to mouse gene IDs identified as human homologs through HCOP.

In addition to the data included in **Table 2**, we plotted the deviation percentages of some graph theoretic measures for a selected number of subgraphs in comparison to Petagraph. As shown in Figure 4, **`HUBMAPSC`** and **`GTEX_EQTL`** exclusion would result in the largest deviation in the number relationships (**Figure 4a**), while **`LINCS`** and **`CMAP`** have the second and third rank in terms of the triangles introduced. However, their contribution is two orders of magnitude smaller than the **`GTEX_COEXP`**, as discussed earlier (**Figure 4b**). Interestingly, the number of label propagation algorithm (LPA) communities introduced by **`GTEX_COEXP`** and **`LINCS`** gave rise to a similar deviation percent to Petagraph but in different directions; while the integration of the **`LINCS`** dataset increased the number of communities by about 0.78%, **`GTEX_COEXP`** decreased this number (**Figure 4c**).

Semantic Types are Petagraph nodes specified to assign types to different entities that are presented as (Concept-Code-Term) triplets to the graph. Currently, there are 127 Semantic Types in Petagraph attached to Concept nodes through **`STY`** relationship types. To understand how the node and relationship data connect such Semantic Types, we picked 54 Semantic Types and listed the relationships (type/SAB) for pairwise combinations of these Semantic Types and presented their quality and quantity as heatmaps (**Figure 5**).

**Figure 5a** shows whether there exists at least one relationship type between at least a pair of Concept nodes of the given start and end Semantic Types. In this regard, this figure can provide intuition into the extent of connectivity in the graph from one point of view, i.e., it shows all pairs of types with no graph-wide relationship between them. Understanding these gaps opens an opportunity to connect the graph across those areas either through graph-encoded information (e.g. by link prediction) or external data sources. Also, this information can be viewed as a guide to develop queries that help to deduce information between Semantic Types with no direct connection between them. For example, Concept nodes with **`STY`** of **`Congenital Abnormality`** and **`Clinical Drug`** have no direct relationship in the graph, while both have relationships to **`Physiologic Function`**. Hence, given the abundance of links of those entities to this intermediate node, queries could be utilized to relate the **`STY`** of **`Congenital Abnormality`** with the **`STY`** of **`Clinical Drug`** (**Figure 5a**).

Similar to the approach above, the log_10_ of the number of relationship types (distinguished by the start and end nodes, relationship type, and SAB) per pair of the 54 Semantic Types was computed and plotted as a heatmap (**Figure 5b**). As illustrated in this figure, the highest diversity in interconnectivity was observed for Semantic Types related to body parts, organs, and tissues, followed by chemicals and biological functions. It was also observed that Concepts with the same or similar Semantic Types (on or surrounding diagonal) have the highest chance of being connected through a diverse set of relationships. On the other hand, off-diagonal Semantic Type pairs are of lower probability to have their associated Concept nodes connected through multiple relationship types.

While **Figure 5b** provides information on the diversity of the relationship types connecting Concept nodes, it does not say much about the number of such relationships. This information is presented in **Figure 5c**. As shown here, on-diagonal nodes are of the highest likelihood be connected with a larger number of links reflected by the log10 of their relationship counts, among which Concept nodes with **`gene or genome`** Semantic Types have at least one order of magnitude larger than the second largest pair, and is mostly composed of **`GTEX_COEXP`** relationships (N_REL>40M). Together with the information in **Figures 5a and 5b**, conclusions could be made about which of the graph’s Semantic Types have the majority of relationships and which information could be extracted from such relationships to elicit information where it is not directly available.

We were interested in how connected human genes were to one another through secondary relationships. As stated in the methods section, the distribution of the shortest path lengths between the graph’s **`Gene`** to **`Gene`** (**`HGNC`**-**`HGNC`**) Concept nodes was estimated using a sample of 256 million pairs of such nodes in a subgraph consisting of Concept nodes (excluding data from **`HUBMAPSC`** source) and the result is illustrated in **Figure 6**. As depicted in the histogram, the probability distribution for shortest lengths ranges from 1 to 10, and the majority of **`HGNC`** Concept nodes (>50%) are connected to each other through only one intermediary node (one hop) thus having a path of length 2, followed by length 3 shortest paths (2 hubs), while direct connectivity and 4-length shortest paths are only a small (<3%) fraction and length >5 paths are relatively negligible.

### Use case: Identifying eQTLs linked to genes associated with a disease or disorder

Diseases and disorders can result from dysregulation of gene expression programs. Genetic changes that can influence gene expression in tissues have been measured in the GTEx project as ‘expression’ quantitative trait loci” or ‘eQTLs.’ An eQTL represents a genetic variant whose allele dosage has been shown to influence a particular gene’s expression profile within a certain tissue. All eQTLs for a particular gene may be interesting to an investigator who is researching genetic effects of that gene’s expression related to a disease. To explore how Petagraph can navigate these kinds of questions, we designed a query that can take a phenotype term and then retrieve all genes associated with that phenotype and then quickly return all tissue-specific eQTLs related to those genes based on the GTEx project. We can also easily restrict the output to a particular tissue, as we did in **Figure 7**, where we queried on MP “Atrial Septal Defects” (MP:0010403) and all phenotype child terms, used the genes associated with that phenotype to map to human genes and and then output all human heart eQTLs associated with the human genes implicated with that phenotype. This query took under fifteen seconds to complete on a laptop computer.

### Use case: What tissues express the most of a certain gene type?

We also utilized Petagraph to show how disparate data types – phenotypes and protein glycosylation – can be quickly linked together for interesting observations. Glycosyltransferases are enzymes that catalyze the transfer of sugar molecules, resulting in the synthesis of saccharides and glycoconjugates. Aberrant gene expression levels of glycosyltransferases have been implicated in multiple neurodegenerative diseases including Alzheimer’s disease, Parkinson’s disease, and Amyloid Lateral Sclerosis (ALS) (Moll et al. 2020; Schneider and Singh 2022). Consequently, it may be interesting to query the expression levels of the glycotransferase gene family in brain tissue compared to other regions of the body. In **Figure 8** we show the median per-tissue expression from GTEx tissues for 221 human glycosyltransferase genes in selected tissues. This may be useful for showing which tissues may be most affected by the loss or aberrant expression of one of these genes. This query links the human glycosyltransferase data set from the GlyGen project and the tissue level expression data set from GTEx. This use case demonstrates the ease in which one can easily query gene expression levels for specific gene identities from other parts of the graph.

### Use case: Linking glycosylation data to models of human diseases and disorders

We also utilized Petagraph to show how disparate data types – phenotypes and protein glycosylation – can be quickly linked together for interesting observations. In this query, we join the major categories of human congenital heart defects (CHDs) – the most common type of structural birth defect – with cellular glycosylation processes as described in the methods section. This type of analysis would be interesting to people studying the relationship between developmental biology, birth defects, and protein post-translational modification. A manual approach to this type of analysis would typically require at least an hour’s worth of work merging across several source files. By using Petagraph, this analysis took seconds.

In this query, predicted glycosylation sites on proteins were extracted by selecting their gene names associated with the selected phenotypes either in mouse or human. As depicted in **Figure 9**, we compared glycosylation processes with three different measures between the CHD-related genes derived from HPO-gene and MP-gene associations (the top and bottom rows, respectively). These measures include the ratio of the genes with glycosylated proteins to all genes (**Figures 9a** **& 9d**), the average number of distinct glycans per glycosylated proteins (**Figures 9b** **& 9e**), and the average number of glycosylation sites/types per glycosylated proteins (**Figures 9c** **& 9f**). Values for the control group of “all proteins” were for `HGNC`-labeled genes without conditioning on an association with any disease. As shown in **Figures 9a** **& 9d**, genes that have an association with CHD have a considerably higher number of protein glycosylation sites compared to all genes in the graph, and that holds for the majority of CHDs, no matter whether `MP` or `HP` is used to link the genes. In **Figure 9a**, heterotaxy (HTX) can be seen to be the only phenotype that is associated with genes with proteins with a lower-than-average ratio of glycosylation sites compared to all genes, while septal defects (atrial (ASD) and ventricular (VSD)) show the highest ratios collectively. Interestingly, the analysis quickly showed that the majority of CHDs have a more diverse set of glycans involved in their glycosylation than all human glycosylated proteins **Figures 9b** **& 9e**. Finally, the average number of glycosylation sites/types per glycosylated proteins are not as deviated from that of all human proteome as the previous measures, however HTX-associated genes retrieved through HPO have a remarkably higher average number of sites/types compared to all glycosylated proteins (>3 folds), that is, even though such genes are not as abundant, their protein products are highly glycosylated (**Figure 9c**).

### Use case: Identifying cell type markers also associated with disorders and diseases

Congenital heart defects are abnormalities in heart structure that occur within the developing human heart early in embryo development (Waters and Hughes 2017; Bennett et al. 2019). By integrating the developmental heart single-cell RNA-seq data into Petagraph, we can now link cell type and gene expression data across other ontologies and datasets. For example, we can take advantage of the gene node’s relationships to other ontologies by filtering the data based on gene location, chromosome, transcript type, expression in tissue (UBERON and GTEx) or biological function (GO). Ultimately, we are able to query Petagraph for DEGs that could be associated with CHDs as defined by the Kids First Data Resource Center in order to identify potential targets for further research. **Figure 10** shows that we can identify cell type clusters that express genes that are linked to a phenotype (Atrial Septal Defects) derived from mouse perturbation study data hosted at the IMPC.

### Use case: Drug side effects and tissues of interest

The drug rofecoxib was recalled due to safety concerns because of an observed increased risk for cardiovascular events, most notably heart attack, and stroke (Topol 2004). **Figure 11** summarizes the result of a query used to extract relationships between a generic drug (rofecoxib) and human tissues. This query is based on the genes with higher than the threshold (TPM>5) expression levels with relationships with rofecoxib either in **`CMAP`** or **`LINCS`** data. An example of such an interconnected set of nodes is given in **Figure 11c**. As shown in Figures 11a and 11b, the ranked tissues point to different organs, i.e. in the case of **`CMAP`** we observed brain tissues more abundantly (e.g. cortex of frontal lobe or hypothalamic structure, **Figure 11a**) while **`LINCS`** provides a correct prediction that heart and blood vessels in the GTEx dataset are most closely related to perturbation gene profiles in rofecoxib (e.g. “right auricular appendage,” “myocardium of left ventricle,” “Coronary arteries,” **Figure 11b**). The **`CMAP`** subgraph represents the CMap pilot study, while the **`L1000`** dataset represents the much larger L1000 project, which tested rofecoxib across 9 cell lines. (Subramanian et al. 2017). **Figure 11d** illustrates how different tissues independently ranked with respect to these datasets correlate to each other, for example, some liver tissues, testis, and blood are predicted to be amongst the least affected by the dug in both datasets, while the structure of the kidney cortex is highly affected as indicated in both datasets.

## Discussion

In this paper we have introduced Petagraph, a biomedical knowledge graph utilizing the base structure of UBKG which is a graph implementation of the UMLS. At the time of publication, we have integrated 15 data sources and mappings into the graph. The majority of these data sources are concerned with genetic and phenotypic data and their cross-mappings between human and mouse (IMPC, MGI, HP and MP, ClinVar, OMIM). We have also ingested primary data sources that contain information about how genes are affected by drugs and chemicals (LINCS L1000, and CMap) as well as gene pathway resources from MSigDB. We now have these datasets integrated into a powerful knowledge graph containing millions of ontological and semantic mappings which can be leveraged to enrich queries and provide contextual and categorical meaning to biomedical questions. By “bringing the data to the ontologies” we have created a large-scale data resource rich in semantic and categorical annotation that allows for rapid and meaningful biomedical queries.

Petagraph’s biomedical data models can easily integrate new ontology structures or experimental datasets utilizing one of many commonly used biomedical terminologies, including HGNC gene IDs, chromosomal locations, small molecule registries, and phenotype ontologies. Petagraph’s base module, UBKG, is updated on a regular basis, and integration of ontologies and datasets is supported by an agnostic data ingestion protocol as discussed.

We have shown that a query in Petagraph can rapidly link across multiple different types of data. For example, we can link diseases or phenotypes to genes, and then those genes can be linked as cell-type markers to single-cell data analyses (**Figure 10**), eQTLs in GTEx (**Figure 7**), or glycotransferases from GlyGen (**Figure 8**). Conversely, collections of gene markers in single-cell data can be linked back to likely phenotypes or pathways through a Petagraph query, such as linking genes with particular glycosylation patterns back phenotypes (**Figure 9**). All of these queries would be of value for those investigators interested in rapid evaluation of large-scale datasets, from simple annotation to more complex evaluation of the data.

Petagraph’s many applications will be useful at every stage of the research process, from hypothesis generation and hypothesis testing to returning data. Petagraph enables users to query and answer non-trivial biomedical questions quickly and easily as we have shown in the Use cases above. Petagraph can also be queried to retrieve relevant data for further downstream analysis and to be used in multi-layer queries where results of one query can be used as a basis for another query. The graph can also be used to intelligently filter queries based on dozens of relevant biomedical criteria.

It would benefit the larger community to expose Petagraph for use to the community through a public API. Currently, Petagraph must be set up in a Neo4j’s community server, which is free but requires use of the Cypher query language. Also, Petagraph is a large, complex graph, and if a user is not familiar with Petagraph’s schema, it could easily return misleading results. This issue can be compounded further if a user is not well-versed in Cypher; although the syntax of the query language is simple, methods like pattern matching and graph traversal can quickly become complex. To remedy these limitations we are building an API so that common types of queries can be parameterized.

Petagraph’s next dataset ingestions will include data from protein-protein interactions, human to mouse developmental stage mappings, genetics testing, multiple -omics datasets and more. Additionally, we plan on developing an API around Petagraph to enable the parameterization of queries so that researchers do not need to understand the graph schema or the query language to take full advantage of this resource.

Petagraph can leverage the rich, semantic, biomedical environment of UMLS in concert with large-scale biomolecular datasets in order to answer more meaningful questions related to human diseases, disorders, drugs and other interesting characteristics. Future implementations of Petagraph will include machine learning analyses to predict important characteristics of genes, diseases and therapeutics.

## Acknowledgements

Funding for this project is acknowledged from the NIH Common Fund, through the Office of Strategic Coordination/Office of the NIH Director under award R03OD030600 (D.M.T); OT2OD026663, OT2OD026675 (J.C.S.); The Department of Biomedical Informatics at The Children’s Hospital of Philadelphia (D.M.T.)

## SUPPLEMENTARY DATA

### Supporting Ontologies, Standards, and Mappings

The following ontologies were added by the UBKG team in support of the Petagraph project:

**Table S1.**

| Abbreviation | Name | Description | Availability |
| --- | --- | --- | --- |
| PATO | Phenotypic Quality Ontology |  |  |
| UBERON | Uber Anatomy Ontology |  |  |
| CL | Cell Ontology |  |  |
| DOID | Human Disease Ontology |  |  |
| CCFASCTB |  |  |  |
| OBI | Ontology for Biomedical Investigations |  |  |
| EDAM |  |  |  |
| HSAPDV | Human Developmental Stages Ontology |  | HsapDv 107 |
| SBO | Systems Biology Ontology |  |  |
| MI |  |  |  |
| CHEBI | CHEBI Integrated Role Ontology |  |  |
| MP | Mammalian Phenotype Ontology |  |  |
| ORDO | Orphanet Rare Disease ontology |  |  |
| UNIPROTKB | UniProtKB |  |  |
| UO | Units of Measure Ontology |  |  |

The UBKG ontology set is regularly updated and can be found at: https://github.com/dbmi-pitt/ubkg/blob/main/scripts/README-PARAMETER%20ORDER%20for%20generation.md

The **mammalian phenotype ontology (MP)** describes the hierarchy of mouse phenotypes. Data was obtained from https://bioportal.bioontology.org/ontologies/MP and directly loaded into an empty neo4j graph instance using the neosemantics tool from neo4j. This graph was then extracted, formatted to fit the UMLS KG schema and loaded into the UMLS KG. The purpose of using the neosemantics (N10s) plugin is that it can render OWL/turtle/triple files into a graph structure. The MP consists of ∼13,200 phenotypes. The hierarchical nature of the MP allows for recursive searches which can be useful when asking broader questions. For example, one can search for all genes related to the heart disease MP term or all genes related to heart disease and all of the child terms of heart disease.

**Human-to-Mouse ortholog mappings** were obtained in April 2021 from the HGNC Comparisons of Orthology Predictions (HCOP) tool at https://www.genenames.org/tools/hcop/. We created new mouse gene Concepts and mapped them using the HCOP data to their corresponding human ortholog. Each orthologous pair share reciprocal relationships, (‘has_human_ortholog’, ‘has_mouse_ortholog’) and out of the 41,638 HGNC Codes in the UMLS, the HCOP tool found at least one mouse ortholog for 20,715 HGNC Codes.

**Mouse genotype-to-phenotype mappings** were obtained from multiple datasets from two databases, namely, the international mouse phenotyping consortium (IMPC) and the mouse genome informatics (MGI) database. The datasets from IMPC and MGI were combined to create a master genotype-to-phenotype dataset. This master dataset contains 10,380 MP terms that are mapped to at least one gene and 17,936 genes that are mapped to at least one MP term.

**Human-to-mouse phenotype mapping** data that connects HPO terms to MP terms was generated using the PheKnowLator tool in December 2020 [PheKnowLator citation.] Here we only map mouse to human phenotypes that are present in the Gabriella Miller Kids First (GMKF) datasets in this instance of Petagraph, to support the use cases in this study, but other mappings could be included at a later date. The mappings that PheKnowLator generated were then checked and edited manually for accuracy. We kept only the highest quality mappings which left us with ∼1000 mappings. Mapping all HPO to MP terms is an ongoing project by the MONDO and uPheno projects [CITE].

**HGNC-to-HPO mappings** data which containing from human genes to human phenotypes were obtained in July 2021 from https://ci.monarchinitiative.org/view/hpo/job/hpo.annotations/lastSuccessfulBuild/artifact/rare-diseases/util/annotation/phenotype_to_genes.txt. These data contain 4,545 genes mapped to at least one phenotype and 10,896 phenotypes mapped to at least one gene.

### Primary Data Sources

#### GlyGen selected datasets

Five datasets from the GlyGen website (https://data.glygen.org) (York et al. 2020)) were chosen based on their relevance to our preliminary use cases. The first two datasets were simply lists of genes that code for glycosyltransferase proteins in the human (https://data.glygen.org/GLY_000004) and mouse (https://data.glygen.org/GLY_000030). These datasets were modeled by creating a human glycosyltransferase’ Concept node as well as a ‘mouse glycosyltransferase’. Then the Concept nodes for the genes (**`HGNC`** nodes) were connected to their respective glycosyltransferase nodes with a ‘is_glycotransferase’ relationship. The next three datasets contain human O-linked and N-linked glycosylation information, namely O-GlcNac (human_proteoform_glycosylation_sites_o_glcnac_mcw.csv v1.12.3), Glyconnect (human_proteoform_glycosylation_sites_glyconnect.csv v1.12.3) and UniCarbKB (human_proteoform_glycosylation_sites_unicarbkb.csv v1.12.3) were obtained from GlyGen. These datasets contain information of human proteoform, i.e. the exact residue on a protein isoform which is glycosylated, type of glycosylation and glycans found to bind that amino acid. To define relationships between human proteins from UniProtKB (**`UNIPROTKB`** concept nodes) (Boutet et al. 2016) and Glycans from the CHEBI resource (Hastings et al. 2016) (as included in **`CHEBI`** data) we introduced an intermediary ontology of gylcosylation sites derived from the information included in the mentioned dataset. In that process, we added 15,101 gylcosylation site nodes to the knowledge graph with 30,198 connections (including reverse relationships) to UniProtKB concept nodes on one hand (type “**`has_site`**” and “**`on_protein`**”, SAB: “**`GlyGen`**”) and (including reverse relationships) connections to glycan **`CHEBI`** nodes with type “**`binds_glycan`**” and “**`binds_protein_site`**” (SAB: “**`GlyGen`**”).

**HuBMAP single-cell RNA-seq** data was obtained in January 2022 from the Human BioMolecular Atlas Program data portal at <u>HuBMAP Data Portal</u>. We included 62 10x Genomics V3 single-cell and single-nucleus RNAseq datasets. Of the 62 studies, 20 are kidney tissue, 15 are spleen tissue, 9 are respiratory system tissue, 9 are large intestine tissue, 5 are lymph node tissue and 4 are thymus tissue. To model each study we used the cluster labels to find the average gene expression per cluster for every gene that had a non-zero expression level.

New HuBMAP concept nodes were created and connected to the corresponding gene node (HGNC) and tissue node (usually UBERON or SNOMED). There are also study concept nodes so that one can search individual studies using the HuBMAP study ID. Furthermore, there are cluster nodes which connect the study nodes to the HuBMAP nodes so that individual clusters can be queried efficiently. So, the general schema of the HuBMAP datasets is study node to cluster node(s) to HuBMAP nodes where each HuBMAP node is connected to its respective gene and tissue node.

**Single cell Fetal heart data** was obtained from Asp et al. 2019 https://pubmed.ncbi.nlm.nih.gov/31835037/ Average gene expression of each cluster was calculated and used to represent each gene within a cell type cluster. Single cell fetal heart concept nodes were created and connections to cell type nodes (author defined cell types, as many Cell Ontology concepts defined in paper are not currently part of the Cell Ontology) and HGNC nodes connections were made.

**IMPC: Mouse genotype-to-phenotype data** were obtained in January 2021 from multiple datasets from two separate databases. The first set of datasets were obtained from the international mouse phenotyping consortium (IMPC), which includes data from KOMP2, and can be found at http://ftp.ebi.ac.uk/pub/databases/impc/all-data-releases/latest/results/. We used the <u>genotype-phenotype-assertions-ALL.csv.gz</u> and the <u>statistical-results-ALL.csv.gz</u> datasets from this database. Both datasets contain, among other data, phenotype to gene mappings in the mouse. The second set of datasets were obtained from the mouse genome informatics (MGI) database and can be found at http://www.informatics.jax.org/downloads/reports/index.html#pheno. We used the <u>MGI_PhenoGenoMP.rpt</u> (Table 5), <u>MGI_GenePheno.rpt</u> (Table 9) and <u>MGI_Geno_DiseaseDO.rpt</u> (Table 10) datasets. All 3 datasets contain, among other data, phenotype-to-gene mappings. The datasets from IMPC and MGI were combined to create a master genotype-to-phenotype dataset. This master dataset contains 10,380 MP terms that are mapped to at least one gene and 17,936 genes that are mapped to at least one MP term.

**Human gene expression vs tissue location data** was obtained in January 2021 from **Genotype-Tissue Expression (GTEx)** Portal (Version 8) at https://gtexportal.org/home/datasets. We used the gene expression dataset, GTEx_Analysis_2017-06-05_v8_RNASeQCv1.1.9_gene_median_tpm, as well as an expression quantitative trait loci (eQTL) dataset, GTEx_Analysis_v8_eQTL. The gene expression dataset contains 54 tissues and 56,200 genes (transcripts). The eQTLs dataset contains over 1.2 million eQTLs from 49 tissues. We created Concept nodes for each eQTL and each tissue-gene expression pair. The eQTL Concepts were then connected to their corresponding tissue node (UBERON), gene node (HGNC) and variant node (dbSNP). The gene expression nodes are connected to their corresponding tissue node and gene node. We also integrated quantitative data from GTEx including p-values for the eQTL data and transcripts per million (TPM) for the gene expression data into the graph. In order to reduce redundancy of nodes we created bin nodes for these quantitative data types. For example, if the gene TP53 has a TPM of 10.5 in the heart, the GTEx Concept for ‘TP53 – Heart’ will be connected to the ’(10,11] ’ TPM bin node. Similarly, if an eQTL has a p-value of 0.0005 it will be connected to the ‘0.0001,0.001’ p-value bin node.

**Kids First phenotypes** were added because we are specifically interested in evaluating patients in the Kids First database. We added phenotypes from 5006 subjects, modeled as Concept nodes, and connected them to their respective HPO Concepts in the graph. KFPheno_June2022_forKG.xlsx, 5006 patient IDs Data containing mappings from human genes to human phenotypes (**`HGNC-HPO`**) were obtained from https://ci.monarchinitiative.org/view/hpo/job/hpo.annotations/lastSuccessfulBuild/artifact/rare-diseases/util/annotation/phenotype_to_genes.txt. These data contain 4,545 genes mapped to at least one phenotype and 10,896 phenotypes mapped to at least one gene.

#### ClinVar

The ClinVar human genetic variants-phenotype submission summary dataset (`2023-01-05`) was utilized to define relationships between human genes and phenotypes (Landrum et al. 2018). For that purpose we only considered genes with pathogenic, likely pathogenic and pathogenic/likely pathogenic variants, and associations with no assertion criteria met were excluded. We also did not include variants that affect a subset of genes (where there was no one-to-one relationship between a gene and phenotype/disease). To retrieve the target phenotype/disease we used MedGen IDs listed in the ClinVar dataset (also already present in the KG). As a result, ClinVar gave rise to 214,040 new relationships (including reverse relationships) with the following characteristics [Type: “**`gene_associated_with_disease_or_phenotype`**”, SAB: “**`CLINVAR`**”] and [type: **`inverse_gene_associated_with_disease_or_phenotype`**, SAB: “**`CLINVAR`**”] connecting **`HGNC`** and **`MEDGEN`**, **`MONDO`**, **`HPO`**, **`EFO`** and **`MESH`** Concept nodes. The ClinVar website is hosted at https://www.ncbi.nlm.nih.gov/clinvar/.

#### MSigDB

MSigDB is a collection of gene set resources, curated or collected from several different sources (Subramanian et al. 2005; Liberzon et al. 2015). Five subsets of MSigDB v7.4 datasets were introduced as entity-gene relationships to the knowledge graph: C1 (positional gene sets), C2 (curated gene sets), C3 (regulatory target gene sets), C8 (cell type signature gene sets) and H (hallmark gene sets). With this subset, we created **`MSIGDB`** Concept nodes for 31,702 MSigDB systematic names (used as Codes) through base64 encoding (Code SAB: **`MSIGDB`**). The relationships between these Concept nodes and **`HGNC`** nodes were established considering the information available in each of the mentioned 5 subsets where the subset information was included in the relationship SABs as “**`MSIGDB`**”. The Term names and SUIs were also compiled according to the MSigDB generic entity names and base64 conversion of such names, respectively. Collectively, MSigDB ingestion added 2,610,854 **`CUI-CUI`** relationships (including reverse relationships) to the KG. Table S2 summarizes the bidirectional relationship information for the MSigDB dataset integration.

**Table S2.**

| MSigDB relationship types and SABs |  |
| --- | --- |
| Relationship Type | Source |
| chr_band_contains_gene | MSigDB C1 |
| inverse_chr_band_contains_gene | MSigDB C1 |
| pathway_associated_with_gene | MSigDB C2 |
| inverse_pathway_associated_with_gene | MSigDB C2 |
| targets_expression_of_gene | MSigDB C3 |
| inverse_targets_expression_of_gene | MSigDB C3 |
| has_marker_gene | MSigDB C8 |
| inverse_has_marker_gene | MSigDB C8 |
| has_signature_gene | MSigDB H |
| inverse_has_signature_gene | MSigDB H |

#### LINCS L1000

We introduced gene-small molecule perturbagen relationships to the KG based on the **`LINCS`** L1000 [iii] edge list available on the Harmonizome database: https://maayanlab.cloud, (Duan et al. 2014; Rouillard et al. 2016). These relationships were summarized from LINCS L1000 CMAP Signatures of Differentially Expressed Genes for Small Molecules dataset. This was done by first finding the corresponding **`CHEBI`** Concept nodes for the L1000 small molecules and then establishing the relationship of such nodes to the KG **`HGNC`** nodes according to the edge list mentioned above. For that purpose, the relationships were collapsed to exclude the cell line, dosage, and treatment time information but the effect directions were retained in relationship types. This led to 3,198,094 relationships (bidirectional) with “**`LINCS`**” as the SAB and types of “**`negatively_correlated_with_gene`**”, “**`positively_correlated_with_gene`**”, “**`inverse_negatively_correlated_with_gene`**” and “**`inverse_positively_correlated_with_gene`**”.

#### Connectivity MAP (CMAP)

In a similar manner to L1000 data integration discussed above, we obtained the edge lists of the CMAP Signatures of Differentially Expressed Genes for Small Molecules dataset from the Harmonizome database :https://maayanlab.cloud, (Lamb et al. 2006; Rouillard et al. 2016). The data was computed based on an earlier study (Lamb et al. 2006; Rouillard et al. 2016). The dataset added 2,625,336 new relationships (including reverse relationships) connecting the KG **`CHEBI`** and **`HGNC`** nodes with types types of “**`negatively_correlated_with_gene`**”, “**`positively_correlated_with_gene`**”, “**`inverse_negatively_correlated_with_gene`**” and “**`inverse_positively_correlated_with_gene`**” and **`SAB`** of “**`CMAP`**”.

#### Human Gene Coexpression

Coexpression of human genes was computed using the GTEx gene TPM (transcript per million) normalized data (GTEx_Analysis_2017-06-05_v8_RNASeQCv1.1.9_gene_tpm by first computing the correlation matrix for each of the GTEx tissues and selecting the entries with pairwise Pearson’s correlation above 0.99. This was done for 54 human tissues and cell types as a result 42,713,122 (including reverse) relationships were ingested into the KG connecting **`HGNC`** Concept nodes (type “**`coexpressed_with_Tissue`**”, SAB: “**`GTEX_COEXP`**”} and (type “**`inverse_coexpressed_with_Tissue`**”, SAB: “**`GTEX_COEXP`**”}.

### Software

A series of python scripts have been developed to ingest specific datasets, each are described on the project GitHub webpage.

https://github.com/TaylorResearchLab/CFDIKG

**Table S3.** Basic Comparison between UBKG and Petagraph.

|  | UBKG | Petagraph | Notes |
| --- | --- | --- | --- |
| **Total # of Nodes** | **19,894,444** | **49,624,263** | **Ex. +50%** |
| **# of total edges** | **52,087,930** | **209,878,717** |  |
| **# of node types** | **5** | **5** | **Concepts, Codes, Terms, Definitions, Semantic** |
| **# of Concepts** | **4,612,422** | **19,000,094** |  |
| **# of Codes** | **5,453,973** | **19,889,354** |  |
| **# of Terms** | **9,803,663** | **10,565,175** |  |
| **# Concept - Concept edges** |  |  |  |
| **# of SAB** |  |  |  |
| **# of edge types** | **1102** | **1911** |  |
| **# of property keys** | **11** | **13** | **Added 'lowerbound' and 'upperbound' properties to numerical bin Concepts** |

**Figure S1:**
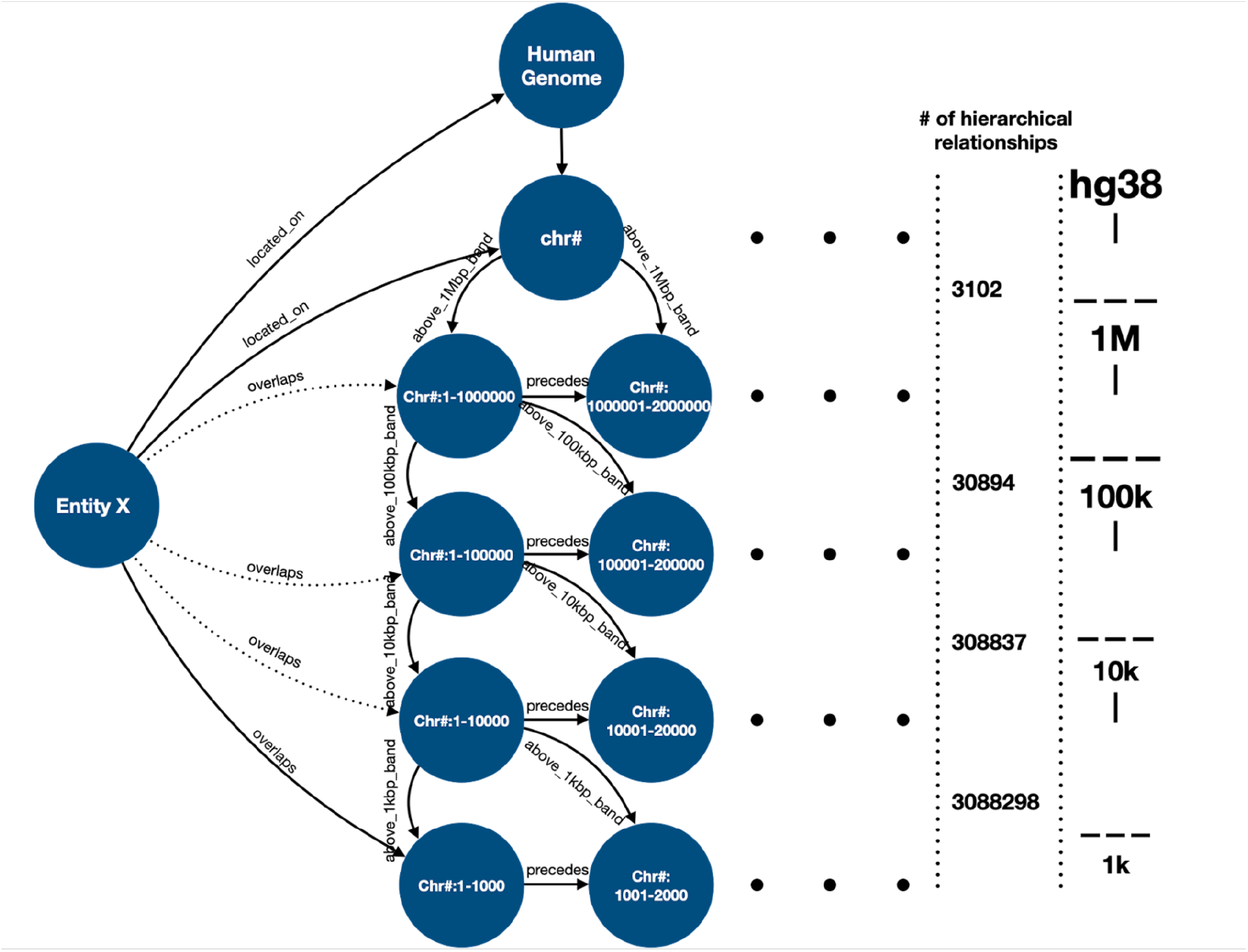
The schema of the Chromosome Region ontology developed for Petagraph. Entity X could be any chromosomal feature, including chromosomal bands, genes, exons, introns, regulatory elements, QTLs, variants, accessible chromatin regions, viral integration sites, human endogenous retroviruses, transposons, tandem repeats, chromosomal contact regions, TADs, telomere, centromeres, and any other type.

**Figure S2:**
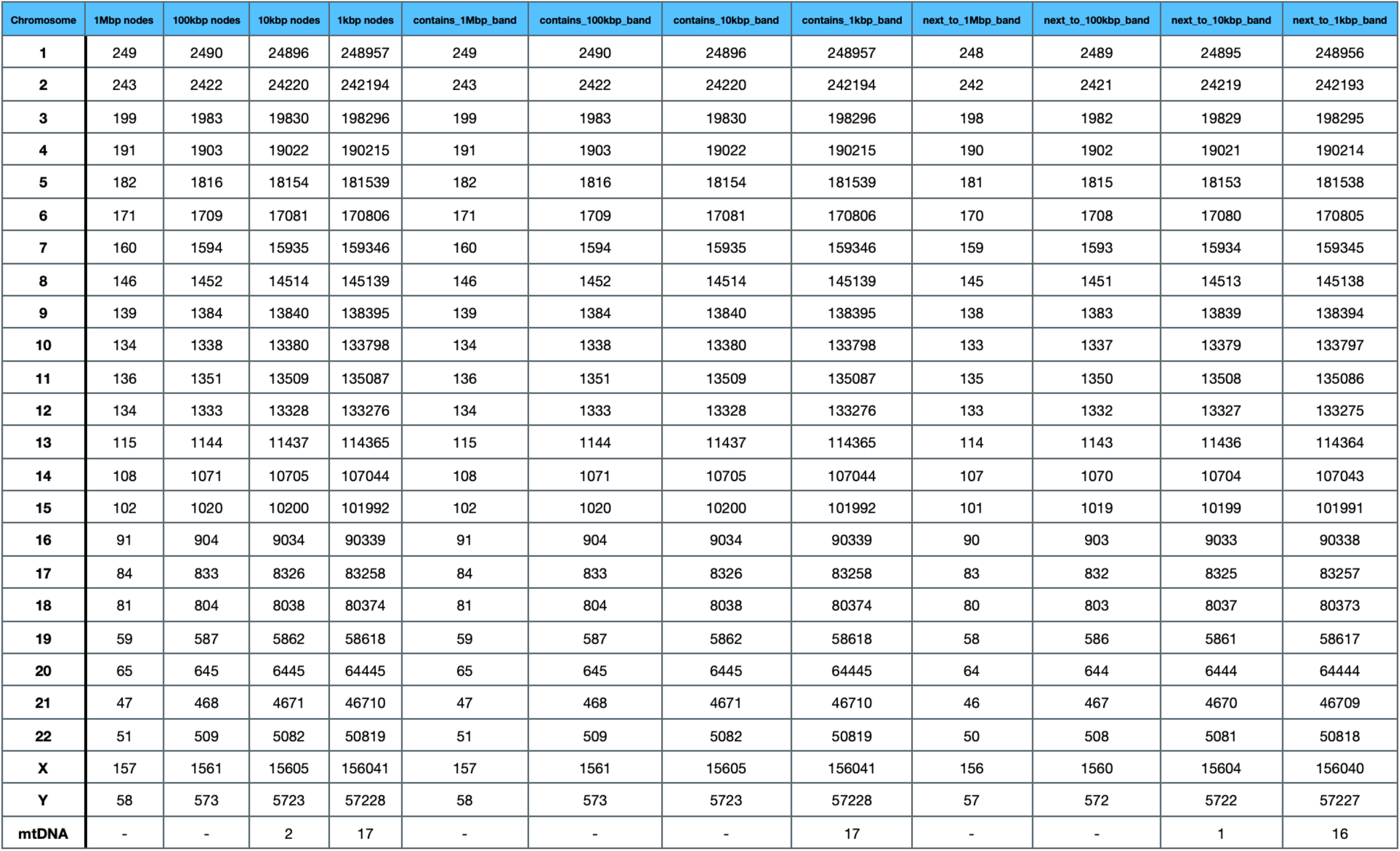
Unidirectional relationship counts between Concepts in the Chromosome Region Ontology

## Notes

https://github.com/TaylorResearchLab/Petagraph

### Competing Interest Statement

The authors have declared no competing interest.

